# Biodistribution of mRNA vaccines in rats: Enrichment in injection site and lymph tissues and rapid clearance without tissue persistence

**DOI:** 10.64898/2026.01.23.701408

**Authors:** Susan M. G. Goody, Christopher Rowbottom, Yun Liu, Nancy Chen

## Abstract

Messenger RNA (mRNA) vaccines using lipid nanoparticles (LNPs) are well-established and globally approved with acceptable safety profiles for preventing respiratory disease. Other mRNA-LNP product concepts are also emerging as novel treatments for broader clinical use. Here, we describe mRNA-LNP vaccine tissue distribution and kinetics after intramuscular dosing using three products formulated with same LNP matrix: mRNA-1273 (Spikevax™), mRNA-1647 (a candidate cytomegalovirus [CMV] vaccine), and a reporter mRNA (nascent peptide-luciferase) drug product. Consistent biodistribution patterns were observed across studies: tissues with highest exposures were the injection site, draining lymph nodes, and spleen, with minimal distribution to non-lymphoid tissues. Vaccine components cleared rapidly from circulation and tissues, with complete elimination simulated to occur by ∼2 weeks. Following mRNA-1273 vaccination, Spike protein levels were transiently observed (elimination <5 days) and did not accumulate with repeated dosing. The ionizable lipid in the LNP matrix, Lipid H, underwent biotransformation and was excreted renally and hepatically, with no human-specific metabolites. Collectively, these results indicate that the LNP composition, not mRNA cargo, governs biodistribution. Furthermore, in a SARS-CoV-2 infection-free model, there was no evidence of Spike protein persistence. Overall, the data establish a framework that justifies leveraging biodistribution data across products and supports eliminating redundant animal studies.

## Introduction

Lipid nanoparticle (LNP)–encapsulated mRNA vaccines are designed for efficient delivery of mRNA to cells adjacent to the intramuscular (IM) injection site and production of a robust immune response.^1^ LNPs typically comprise a mixture of 4 components: an ionizable lipid, a PEGylated lipid, cholesterol, and a helper lipid such as distearoylphosphatidylcholine (DSPC).^2^ ^3^ Following IM injection, mRNA-LNP vaccines are taken up predominantly by immune cells, such as antigen-presenting cells (APCs) and fibroblasts. at the injection site and draining lymph nodes.^4^ APCs expressing the vaccine antigen, along with mRNA-LNP, traffic to afferent lymphatics and initiate seeding of germinal centers in draining lymph nodes, where germinal center reactions drive affinity maturation of antigen-specific B-cell receptors and produce corresponding long-lived plasma and memory B-cell antibodies.^3, 5^

mRNA-LNP vaccines are recognized worldwide for delivering clinically meaningful outcomes with well-characterized and acceptable safety profiles. Global licensure of 4 mRNA-LNP vaccines, including Moderna’s mRNA-1273 (Spikevax™), mRNA-1345 (mRESVIA™), mRNA-1283 (mNEXSPIKE™) and Pfizer–BioNTech’s BNT162b2 (Comirnaty™), are a testament to this.^6–8^ Beyond these licensed products, there are several examples of promising clinical data with other indications such as seasonal influenza^9, 10^ and oncology.^11, 12^ Collectively, these successes solidify mRNA-LNP vaccines and therapies as a versatile, regulatory-validated modality with a growing clinical footprint spanning licensed respiratory vaccines, late-stage prevention therapeutics, and personalized oncology medicines.

Drug developers follow internationally recognized guidelines for preclinical evaluations of mRNA-LNP vaccines,^13, 14^ which provide recommendations on the studies designed to support their regulatory approval. These studies rigorously assess the preclinical immunogenicity, safety, metabolism, clearance, and biodistribution of the vaccines to support testing in humans. Biodistribution studies in rats are an accepted regulatory standard for mRNA-LNP vaccines,^15–19^ and are designed to map the tissue localization of drug components and/or expressed antigen, measure their persistence and clearance, and to assess for potential accumulation with repeated administration if more than one dosing occasion is employed. For vaccines, recommendations for biodistribution studies are risk-based and product-specific rather than prescriptive, noting that bridging across products is acceptable when scientifically justified if LNP composition and route of administration remain unchanged.^13, 14^ Detailed biodistribution datasets have been reviewed and deemed acceptable to support global licensure of mRNA-LNP vaccines; however, because these datasets are proprietary in regulatory submission dossiers, the broader scientific community has limited visibility to the details of the methods used and the results from these studies.

To increase transparency of these data, we present a comprehensive description of the tissue distribution and systemic kinetics of 3 mRNA-LNP products formulated in the same LNP matrix following IM dosing in Sprague Dawley rats. We also provide the first characterization of SARS-CoV-2 Spike protein tissue biodistribution and clearance in a SARS-CoV-2 infection-naïve model, addressing a key limitation of human studies where pre-existing or subclinical SARS-CoV-2 infection cannot be reliably ruled out. Additionally, we describe the metabolism and/or excretion of the ionizable lipid used in the LNP formulations, Lipid H. Altogether, the data substantiate that it is the LNP composition, not mRNA cargo or immune response, that drive tissues distribution and kinetics, and that LNP components and Spike protein levels were transient, with no evidence of persistence beyond 1-2 weeks after IM injection.

## Results

### Across drug products, tissues with the highest exposure of mRNA and Lipid H are consistently the injection site, draining lymph nodes, and spleen

An evaluation of the area under the time–concentration curve (AUC; h*ng/mL or h*ng/g for plasma/serum or tissue, respectively) for mRNA and Lipid H across drug products revealed that the tissues with the highest AUC were consistently the injection site, draining lymph nodes (particularly axillary, inguinal, and/or popliteal), and spleen (Tables 1 and 2). Differences in dose levels and/or final collection timepoints have the potential to introduce variability in the value of raw calculated AUC; therefore, tissue exposures were normalized to systemic exposure (based on AUC). The resulting values (otherwise referred to as the “tissue-to-systemic ratio. expressed as a percentage [%]) were also used to rank order tissue exposure relative to systemic exposure. Tissues with ratios >100% represent those with higher exposure relative to systemic exposure, whereas tissues with ratios <100% represent tissues with lower exposure compared with systemic exposure. The resulting distribution patterns of tissue to systemic ratios were consistent with the calculated AUC value (Tables 1 and 2) across the drug products, as demonstrated by ratios of >100% for lymphoid tissues and <100% for other distal tissues

**Table 1.**
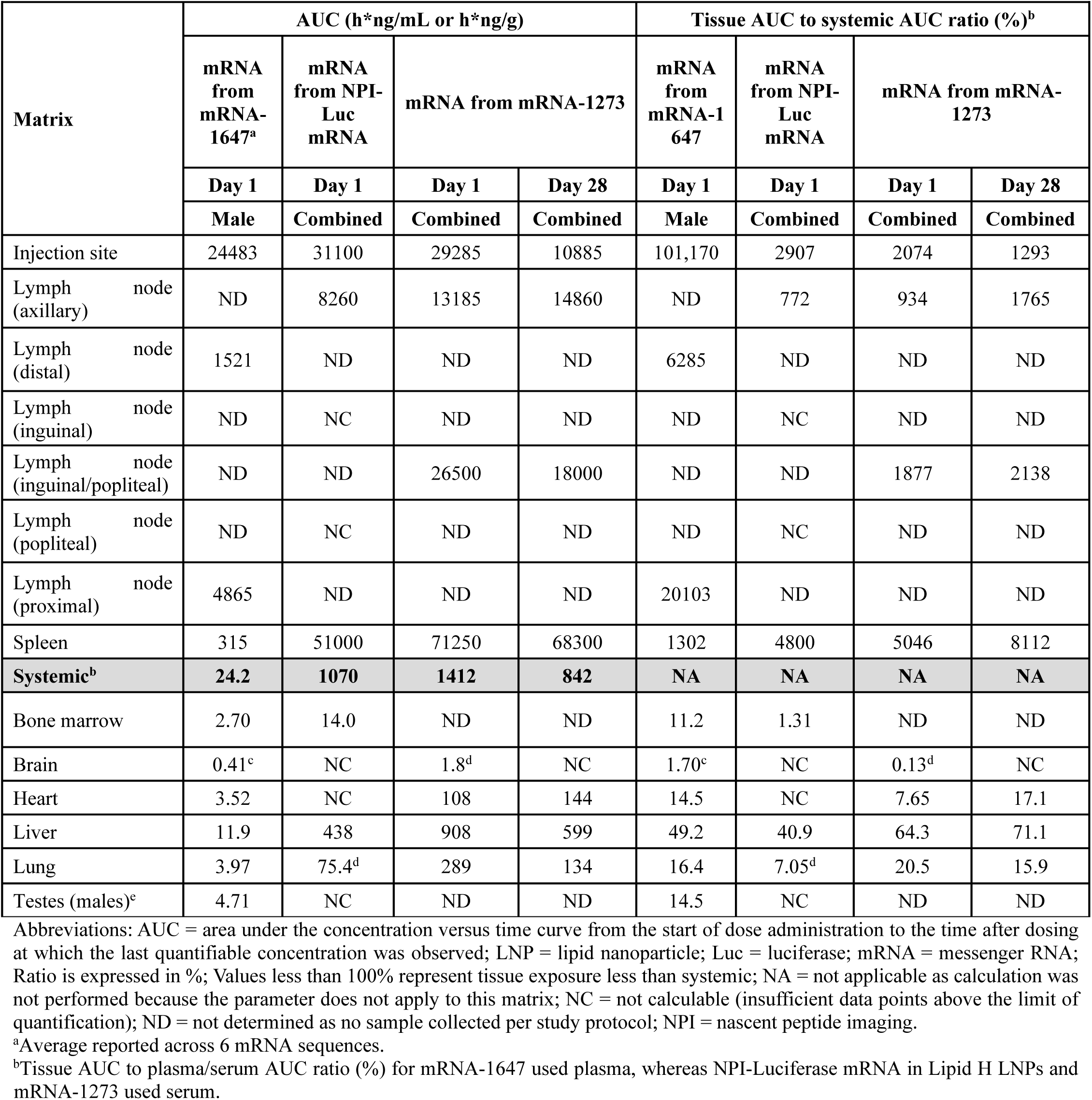
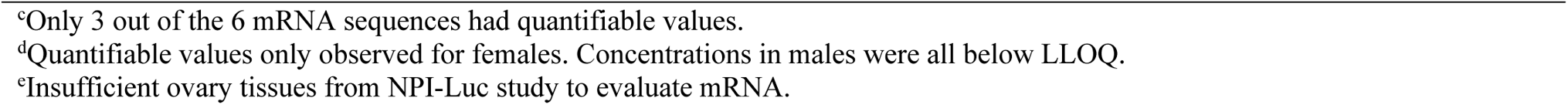
Summary of systemic and tissue mRNA AUC and tissue to systemic AUC ratios (sex combined)

| Matrix | AUC (h*ng/mL or h*ng/g) |  |  |  | Tissue AUC to systemic AUC ratio (%)<sup>b</sup> |  |  |  |
| --- | --- | --- | --- | --- | --- | --- | --- | --- |
|  | mRNA from mRNA-1647<sup>a</sup> | mRNA from NPI-Luc mRNA | mRNA from mRNA-1273 |  | mRNA from mRNA-1647 | mRNA from NPI-Luc mRNA | mRNA from mRNA-1273 |  |
|  | Day 1 | Day 1 | Day 1 | Day 28 | Day 1 | Day 1 | Day 1 | Day 28 |
|  | Male | Combined | Combined | Combined | Male | Combined | Combined | Combined |
| Injection site | 24483 | 31100 | 29285 | 10885 | 101,170 | 2907 | 2074 | 1293 |
| Lymph node (axillary) | ND | 8260 | 13185 | 14860 | ND | 772 | 934 | 1765 |
| Lymph node (distal) | 1521 | ND | ND | ND | 6285 | ND | ND | ND |
| Lymph node (inguinal) | ND | NC | ND | ND | ND | NC | ND | ND |
| Lymph node (inguinal/popliteal) | ND | ND | 26500 | 18000 | ND | ND | 1877 | 2138 |
| Lymph node (popliteal) | ND | NC | ND | ND | ND | NC | ND | ND |
| Lymph node (proximal) | 4865 | ND | ND | ND | 20103 | ND | ND | ND |
| Spleen | 315 | 51000 | 71250 | 68300 | 1302 | 4800 | 5046 | 8112 |
| **Systemic<sup>b</sup>** | **24.2** | **1070** | **1412** | **842** | **NA** | **NA** | **NA** | **NA** |
| Bone marrow | 2.70 | 14.0 | ND | ND | 11.2 | 1.31 | ND | ND |
| Brain | 0.41<sup>c</sup> | NC | 1.8<sup>d</sup> | NC | 1.70<sup>c</sup> | NC | 0.13<sup>d</sup> | NC |
| Heart | 3.52 | NC | 108 | 144 | 14.5 | NC | 7.65 | 17.1 |
| Liver | 11.9 | 438 | 908 | 599 | 49.2 | 40.9 | 64.3 | 71.1 |
| Lung | 3.97 | 75.4<sup>d</sup> | 289 | 134 | 16.4 | 7.05<sup>d</sup> | 20.5 | 15.9 |
| Testes (males)<sup>c</sup> | 4.71 | NC | ND | ND | 14.5 | NC | ND | ND |

Abbreviations: AUC = area under the concentration versus time curve from the start of dose administration to the time after dosing at which the last quantifiable concentration was observed; LNP = lipid nanoparticle; Luc = luciferase; mRNA = messenger RNA; Ratio is expressed in %; Values less than 100% represent tissue exposure less than systemic; NA = not applicable as calculation was not performed because the parameter does not apply to this matrix; NC = not calculable (insufficient data points above the limit of quantification); ND = not determined as no sample collected per study protocol; NPI = nascent peptide imaging.
<sup>a</sup>Average reported across 6 mRNA sequences.
<sup>b</sup>Tissue AUC to plasma/serum AUC ratio (%) for mRNA-1647 used plasma, whereas NPI-Luciferase mRNA in Lipid H LNPs and mRNA-1273 used serum.
<sup>c</sup>Only 3 out of the 6 mRNA sequences had quantifiable values.
<sup>d</sup>Quantifiable values only observed for females. Concentrations in males were all below LLOQ.
<sup>e</sup>Insufficient ovary tissues from NPI-Luc study to evaluate mRNA.

**Table 2.** Summary of systemic and tissue Lipid H AUC and AUC to systemic AUC ratios (sex combined)

| Matrix<sup>a</sup> | AUC (h*ng/mL or h*ng/g) |  |  | Tissue AUC to systemic AUC ratio (%) |  |  |
| --- | --- | --- | --- | --- | --- | --- |
|  | NPI-Luc mRNA | mRNA-1273 |  | NPI-Luc mRNA | mRNA-1273 |  |
|  | Day 1 | Day 1 | Day 28 | Day 1 | Day 1 | Day 28 |
|  | Combined | Combined | Combined | Combined | Combined | Combined |
| Injection site | 152000 | 168750 | 306000 | 2165 | 1023 | 2045 |
| Lymph node (axillary) | 13700 | 54950 | 89100 | 195 | 333 | 595 |
| Lymph node (inguinal) | 83100 | ND | ND | 1184 | ND | ND |
| Lymph node (inguinal/popliteal) | ND | 126500 | 90350 | ND | 767 | 604 |
| Lymph node (popliteal) | 71200 | ND | ND | 1014 | ND | ND |
| Spleen | 6980 | 65450 | 35950 | 99.4 | 397 | 240 |
| **Systemic** | **7020** | **16495** | **14965** | **NA** | **NA** | **NA** |
| Bone marrow | 591 | ND | ND | 8.42 | ND | ND |
| Brain | NC | NC | NC | NC | NC | NC |
| Heart | NC | 21.7 | 28.2 | NC | 0.132 | 0.188 |
| Kidney (left) | 8.09 | ND | ND | 0.115 | ND | ND |
| Kidney (right) | 14.6 | ND | ND | 0.208 | ND | ND |
| Liver | 2180 | 1865 | 3990 | 31.1 | 11.3 | 26.7 |
| Lung | 231 | 366 | 238 | 3.29 | 2.22 | 1.59 |
| Ovaries (females) | 457 | ND | ND | 6.51 | ND | ND |

Abbreviations: AUC = area under the concentration versus time curve from the start of dose administration to the time after dosing at which the last quantifiable concentration was observed; LNP = lipid nanoparticle; Luc = luciferase; mRNA = messenger RNA; Ratio is expressed in %; Values less than 100% represent tissue exposure less than systemic; NA = not applicable as calculation was not performed because the parameter does not apply to this matrix; NC = not calculable (insufficient data points above the lower limit of quantification); ND = not determined as no samples were collected per study protocol; NPI = nascent peptide imaging.
<sup>a</sup>Concentrations in testis were all below LLOQ in NPI-Luc study and were not determined in mRNA-1273 study.

Of the distal tissues, the liver had the highest tissue-to-systemic ratios for mRNA and Lipid H. When compared to the injection site, which serves as the primary depot of exposure after IM administration (Tables 1 and 2), liver exposure was significantly lower, representing only ∼3% relative to the injection site. Other distal tissues, such as bone marrow, brain, heart, lung, testes, or ovaries, had exposures that were <1% relative to the injection site (Tables 1 and 2). The concentrations in these tissues were noted to be highly variable and were either near the lower limit of assay quantification (LLOQ; Tables 8; Table 1and 2) or had measurable mRNA in the absence of lipid. For example, the detectable concentrations in brain were sporadic and inconsistent across sexes (e.g., only in females and at a few early timepoints) and mRNA concentrations were observed near the lower end of the assay limit, and sometimes without quantifiable levels of Lipid H (Table 1 and 2). Collectively, these findings demonstrate that exposures in the brain and other distal tissues are negligible and show limited biodistribution of mRNA-LNP components beyond the injection site and lymphoid tissues.

### No persistence of mRNA, Lipid H, or SARS-CoV-2 Spike protein in the systemic circulation or tissues

In the systemic circulation, mRNA and Lipid H declined rapidly, with a serum half-life (T_1/2_) of approximately 3 h and 8 h, respectively (Table 3). The T_1/2_ values of mRNA in lymphoid tissues with highest exposure (based on AUC) ranged from 24 h to 48 h in lymph nodes and was ∼63 h in the spleen (Table 3). Spike protein concentration peaked at 24 h (the time after dosing at which the maximum observable pharmacodynamic effect occurred [TE_max_]) following mRNA-1273 administration and declined rapidly with an effective serum T_1/2_ of 18 h (Table 4). The quantifiable levels in serum were observed only up to 3 days after the first dose and up to 7 days after the second (Table 4). While there were enough timepoints with quantifiable Spike protein levels in the serum after the first dose to calculate the area under the pharmacodynamic (PD) effect (encoded protein)-time curve from time 0 to the last quantifiable concentration post dose (AUEC_last_), there were not enough timepoints with quantifiable levels after the second dose to calculate the AUEC_last_. Following the same distribution pattern as mRNA and Lipid H, Spike protein was detected mainly in the injection site, spleen and lymph nodes. Liver was the only distal tissue with measurable Spike protein; however, values were near the LLOQ and were observed only up to 24 h. Spike protein was not detected in brain or heart or lung (Table 4).

**Table 3.**
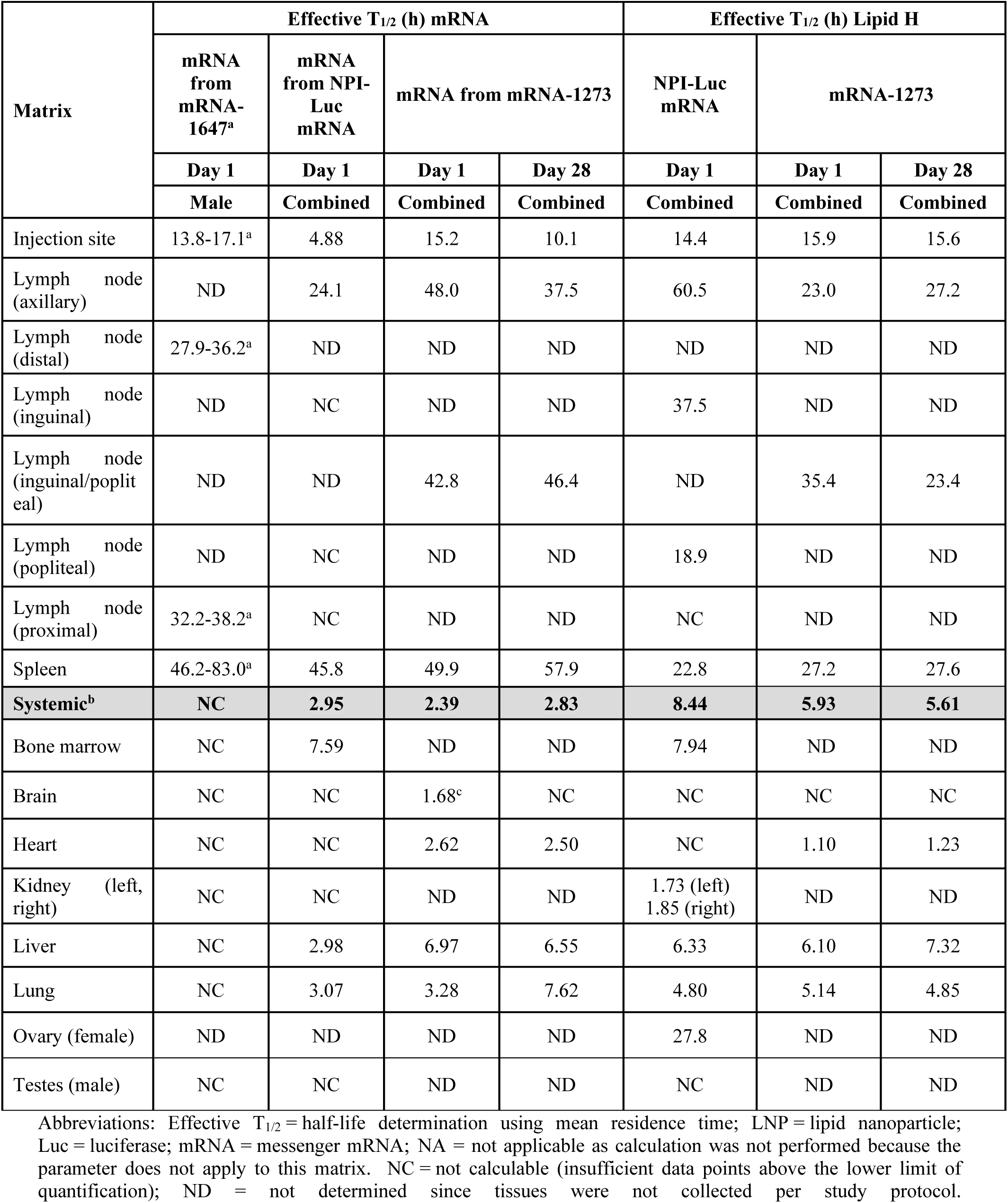
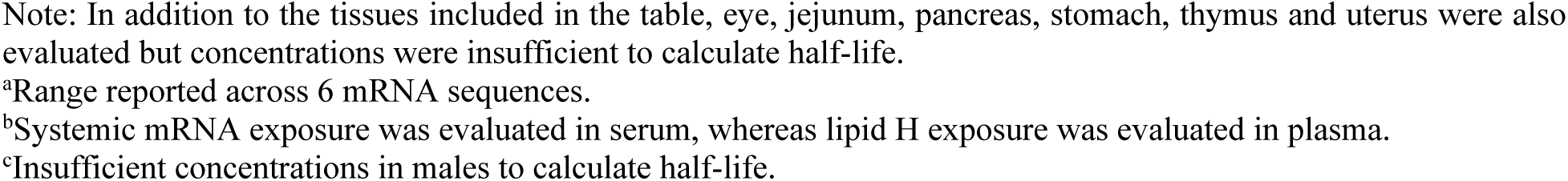
Summary of systemic and tissue mRNA or Lipid H effective T_1/2_ (sex combined)

| Matrix | Effective T<sub>1/2</sub> (h) mRNA |  |  |  | Effective T<sub>1/2</sub> (h) Lipid H |  |  |
| --- | --- | --- | --- | --- | --- | --- | --- |
|  | mRNA from mRNA-1647<sup>a</sup> | mRNA from NPI-Luc mRNA | mRNA from mRNA-1273 |  | NPI-Luc mRNA | mRNA-1273 |  |
|  | Day 1 | Day 1 | Day 1 | Day 28 | Day 1 | Day 1 | Day 28 |
|  | Male | Combined | Combined | Combined | Combined | Combined | Combined |
| Injection site | 13.8-17.1<sup>a</sup> | 4.88 | 15.2 | 10.1 | 14.4 | 15.9 | 15.6 |
| Lymph node (axillary) | ND | 24.1 | 48.0 | 37.5 | 60.5 | 23.0 | 27.2 |
| Lymph node (distal) | 27.9-36.2<sup>a</sup> | ND | ND | ND | ND | ND | ND |
| Lymph node (inguinal) | ND | NC | ND | ND | 37.5 | ND | ND |
| Lymph node (inguinal/popliteal) | ND | ND | 42.8 | 46.4 | ND | 35.4 | 23.4 |
| Lymph node (popliteal) | ND | NC | ND | ND | 18.9 | ND | ND |
| Lymph node (proximal) | 32.2-38.2<sup>a</sup> | NC | ND | ND | NC | ND | ND |
| Spleen | 46.2-83.0<sup>a</sup> | 45.8 | 49.9 | 57.9 | 22.8 | 27.2 | 27.6 |
| **Systemic<sup>b</sup>** | **NC** | **2.95** | **2.39** | **2.83** | **8.44** | **5.93** | **5.61** |
| Bone marrow | NC | 7.59 | ND | ND | 7.94 | ND | ND |
| Brain | NC | NC | 1.68<sup>c</sup> | NC | NC | NC | NC |
| Heart | NC | NC | 2.62 | 2.50 | NC | 1.10 | 1.23 |
| Kidney (left, right) | NC | NC | ND | ND | 1.73 (left)<br>1.85 (right) | ND | ND |
| Liver | NC | 2.98 | 6.97 | 6.55 | 6.33 | 6.10 | 7.32 |
| Lung | NC | 3.07 | 3.28 | 7.62 | 4.80 | 5.14 | 4.85 |
| Ovary (female) | ND | ND | ND | ND | 27.8 | ND | ND |
| Testes (male) | NC | NC | ND | ND | NC | ND | ND |

Abbreviations: Effective T<sub>1/2</sub> = half-life determination using mean residence time; LNP = lipid nanoparticle; Luc = luciferase; mRNA = messenger mRNA; NA = not applicable as calculation was not performed because the parameter does not apply to this matrix. NC = not calculable (insufficient data points above the lower limit of quantification); ND = not determined since tissues were not collected per study protocol.
Note: In addition to the tissues included in the table, eye, jejunum, pancreas, stomach, thymus and uterus were also evaluated but concentrations were insufficient to calculate half-life. <sup>a</sup>Range reported across 6 mRNA sequences. <sup>b</sup>Systemic mRNA exposure was evaluated in serum, whereas lipid H exposure was evaluated in plasma. <sup>c</sup>Insufficient concentrations in males to calculate half-life.

**Table 4.** Summary of SARS-CoV-2 Spike protein quantification and calculated pharmacodynamics (PD) parameters following one or two administrations of mRNA-1273.

| Matrix | Detection<br>(TE<sub>max</sub>/TE<sub>last</sub>)<br>(Day 1 vs. Day 28) | Quantification<br>(E<sub>max</sub>/AUEC<sub>last</sub>) |  |  | Effective T<sub>1/2</sub><br>(if calculable) | Persistence |
| --- | --- | --- | --- | --- | --- | --- |
| Serum | Day 1: TE<sub>max</sub> at 24 h, TE<sub>last</sub> at 72 h<br>Day 28: TE<sub>max</sub> at 168 h, TE<sub>last</sub> at 168 h |  | Day 1 | Day 28 | ~18 h (Day 1 only) | No persistence |
|  |  | E<sub>max</sub><br>(ng/mL) | 5.38 | 1.80 |  |  |
|  |  | AUEC<sub>last</sub><br>(h*ng/mL) | 190 | NC |  |  |
| Liver | Day 1: TE<sub>max</sub> at 10 h, TE<sub>last</sub> at 24 h<br>Day 28: TE<sub>max</sub> at 24 h, TE<sub>last</sub> at 24 h |  | Day 1 | Day 28 | ~4–5 h (Day 1 only) | No persistence |
|  |  | E<sub>max</sub> (ng/g) | 54 | 15 |  |  |
|  |  | AUEC<sub>last</sub><br>(h*ng/g) | 789 | NC |  |  |
| Brain | Not detected at either timepoint | NC |  |  | NC | NC |
| Heart | Not detected at either timepoint | NC |  |  | NC | NC |
| Lung | Day 1: Not detected<br>Day 28: TE<sub>max</sub> at 10 h, TE<sub>last</sub> at 10 h | NC |  |  | NC | No persistence |
| Lymph Nodes Pooled | Day 1: TE<sub>max</sub> at 14 h, TE<sub>last</sub> at 17 h<br>Day 28: Not detected |  | Day 1 | Day 28 | NC | No persistence |
|  |  | E<sub>max</sub> (ng/g) | 341 | NC |  |  |
|  |  | AUEC<sub>last</sub><br>(h*ng/g) | NC | NC |  |  |
| Injection site | Day 1: TE<sub>max</sub> at 17 h, TE<sub>last</sub> at 48 h<br>Day 28: TE<sub>max</sub> at 10 h, TE<sub>last</sub> at 10 h |  | Day 1 | Day 28 | ~7 h (Day 1 only) | No persistence |
|  |  | E<sub>max</sub> (ng/g) | 288 | 29.8 |  |  |
|  |  | AUEC<sub>last</sub><br>(h*ng/g) | 4075 | NC |  |  |
| Spleen | Day 1: TE<sub>max</sub> at 10 h, TE<sub>last</sub> at 24 h<br>Day 28: TE<sub>max</sub> at 10 h, TE<sub>last</sub> at 10 h |  | Day 1 | Day 28 | ~5 h (Day 1 only) | No persistence |
|  |  | E<sub>max</sub> (ng/g) | 77 | 82 |  |  |
|  |  | AUEC<sub>last</sub> (h*ng/g) | 1350 | NC |  |  |

Abbreviations: E<sub>max</sub> = Maximum observed pharmacodynamic effect (encoded protein); AUEC<sub>last</sub> = the area under the pharmacodynamic effect (encoded protein)-time curve from time 0 to the last quantifiable concentration post dose; NC = not calculated (insufficient data points above the lower limit of quantification); TE<sub>last</sub> = the time after dosing at which the last observable pharmacodynamic effect (encoded protein) occurred.

While these T_1/2_ values provide substantive evidence of rapid clearance and lack of persistence, the differences in timing for the final sample collection meant that it was not possible to precisely define the timepoint at which mRNA, Lipid H and/or encoded Spike protein were completely eliminated. Thus, first-order elimination (using observed maximum concentration [C_max_] and serum/tissue T_1/2_ for mRNA, and TE_max_ and effective T_1/2_ for Spike protein) was used to simulate the timepoints at which there was near 100% elimination (Tables 5 and 6; Figure 1A and 1B). To estimate the most extreme circumstances, clearance for tissues with the highest exposure and corresponding T_1/2_ (spleen and lymph nodes) was simulated using the most sensitive analyte (mRNA). The simulation revealed >98% elimination of mRNA at approximately 2 weeks post-dose (Table 5; Figure 1A). In the case of Spike protein in the systemic circulation, the simulation revealed >98% elimination within 5 days post-dose (Table 6; Figure 1B).

**Figure 1.**
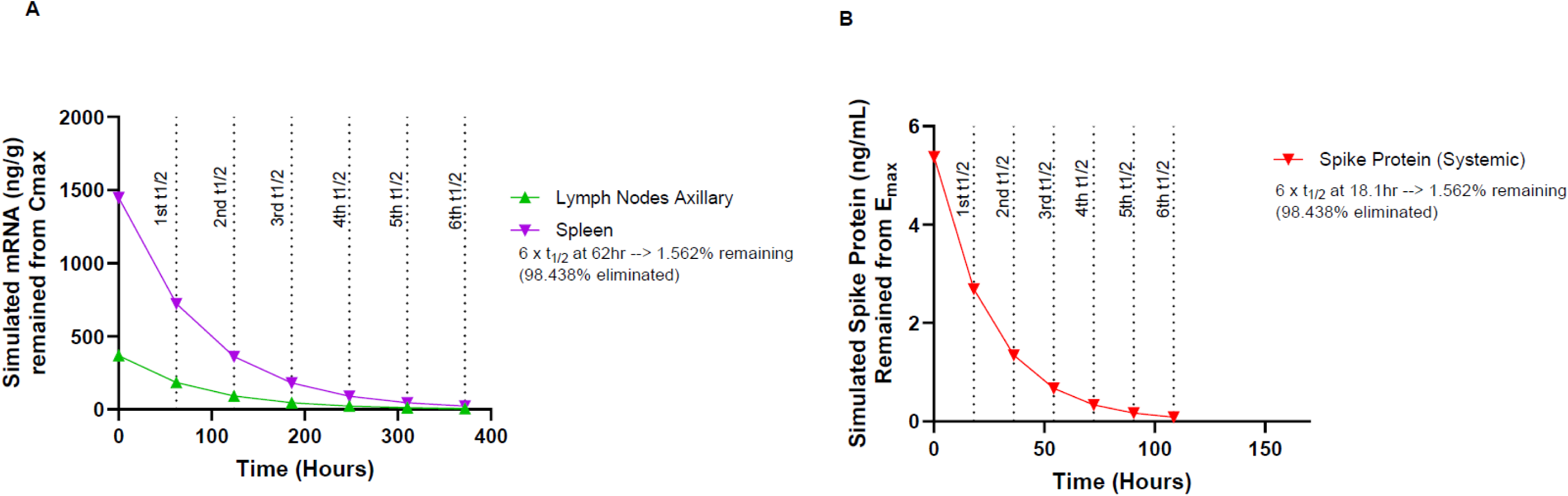
Simulated concentration–time profiles illustrating elimination of mRNA and Spike protein. (A) Simulated mRNA remaining from C_max_ in axillary lymph nodes and spleen. (B) Simulated spike protein remaining from E_max_ in plasma. Profiles were simulated using observed E_max_ and estimated effective half-life (T_1/2_). Vertical dashed lines indicate sequential half-lives. At six half-lives, ∼1.6% of the initial concentration remains (∼98.4% eliminated).

**Table 5.** Simulated mRNA exposures in tissues with the highest AUC and longest T_1/2_ demonstrate near complete elimination approximately 2 weeks after administration of mRNA-1273.

| Half-lives (n) | Time (h) | Simulated mRNA (ng/g) | | Simulated %<br>Eliminated Relative<br>to $C_{max}$ |
| --- | --- | --- | --- | --- |
| | | Remaining from $C_{max}$ | | |
|  |  | Spleen | Lymph Nodes<br>Axillary |  |
| 0 | 0 | 1447.5 | 366.8 | 0 |
| 1 | 62 | 723.5 | 183.4 | 50 |
| 2 | 124 | 361.75 | 91.7 | 75 |
| 3 | 186 | 180.9 | 45.9 | 87.5 |
| 4 | 248 | 90.4 | 22.9 | 93.8 |
| 5 | 310 | 45.2 | 11.5 | 96.9 |
| 6 | 372 | 22.6 | 5.75 | 98.4 |

**Table 6.** Simulated exposures of SARS-CoV-2 Spike protein in systemic circulation demonstrate near complete elimination within 5 days after administration of mRNA-1273.

| Half-lives (n) | Time (h) | Simulated Spike Protein<br>(ng/mL) Remaining from E<sub>max</sub> | Simulated % of<br>Eliminated Relative to<br>E<sub>max</sub> |
| --- | --- | --- | --- |
| 0 | 0 | 5.38 | 0 |
| 1 | 18.1 | 2.69 | 50 |
| 2 | 36.2 | 1.35 | 75 |
| 3 | 54.3 | 0.673 | 87.5 |
| 4 | 72.4 | 0.336 | 93.8 |
| 5 | 90.5 | 0.168 | 96.9 |
| 6 | 108.7 | 0.084 | 98.4 |

The expressed nascent peptide-luciferase protein (NPI-Luc) from the study with NPI-Luc mRNA in Lipid-H-containing LNPs was largely not quantifiable in either tissues or systemic exposure (data not shown).

### Lack of accumulation of mRNA, Lipid H and Spike protein with repeated dosing and no impact of immune response on tissue distribution

To further interrogate the potential for persistence of mRNA and Lipid H in the systemic circulation or in tissues, and to assess whether development of an immune response impacted the biodistribution profile, exposures of mRNA and Lipid H were evaluated following a second dose of mRNA-1273 delivered approximately 4 weeks after the first dose. The AUC and C_max_ of mRNA and Lipid H in the systemic circulation and in tissues with highest exposure (injection site, lymph node, spleen) were the same on Day 1 as on Day 28, with no statistically significant differences observed (Figure 2). Similarly for Spike protein in systemic or tissues, maximum observed PD effect (E_max_) values on Day 1 and Day 28 were either comparable or, in some cases, lower or could not be determined due to insufficient measurable response on Day 28 (Table 4). Comparison of AUEC_last_ between Day 1 and Day 28 was not feasible, as only a few quantifiable levels were available following the second dose to obtain reliable estimation.

**Figure 2.**
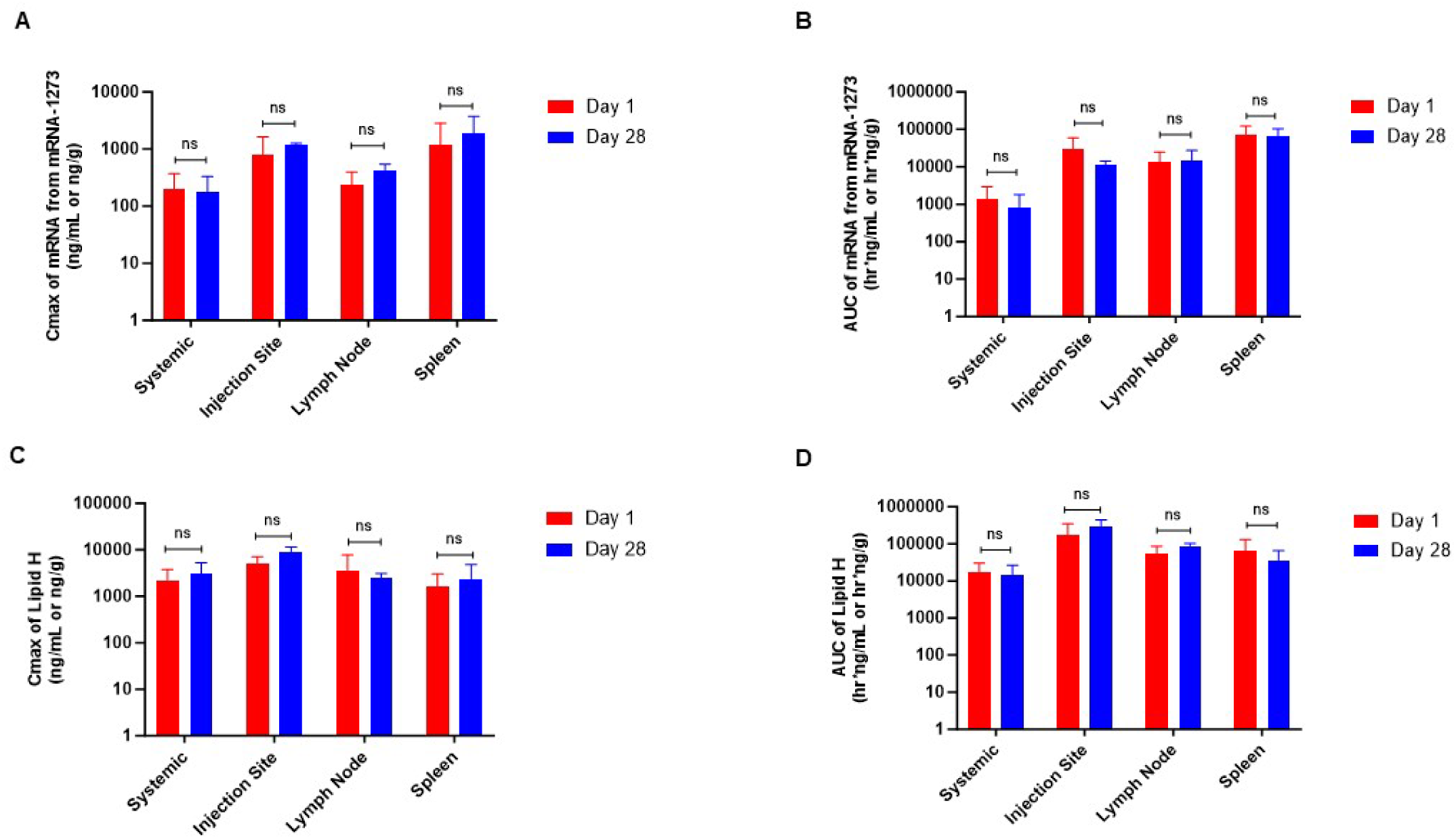
Cmax and AUC of mRNA and Lipid H in Sprague–Dawley rats following intramuscular administration of mRNA-1273 on Day 1 and Day 28. (A) C_max_ of mRNA-1273. (B) AUC of mRNA-1273. (C) C_max_ of Lipid H. (D) AUC of Lipid H. Tissues analyzed included serum, injection site, axillary lymph nodes, and spleen. Bars represent mean ± SD. Comparisons between Day 1 and Day 28 were performed using unpaired t-tests; no statistically significant differences were observed (*p* < 0.05). ns = not significant.

To confirm development of an immune response to mRNA-1273 and further reiterate that immunogenicity does not impact the biodistribution profile, antibody responses against the SARS-CoV-2 Spike protein S2 subunit (S2P) antigen were measured following the first and second dose of mRNA-1273. At baseline (Day 1, pre-dose), antibody titers were at or near the LLOQ (<30 antibody units/mL), with no measurable pre-existing antibodies (Figure 3A). At 2 weeks (336 h) post-first-dose, all animals demonstrated a robust antibody response, with binding titers increasing by several orders of magnitude compared with pre-dose values. On Day 28, prior to the second dose, antibody titers remained elevated relative to baseline, demonstrating persistence of the immune response from the primary immunization (Figure 3B). Following the second dose, titers were further boosted, reaching high and consistent levels across all animals by 2 weeks after the second dose (336 h).

**Figure 3.**
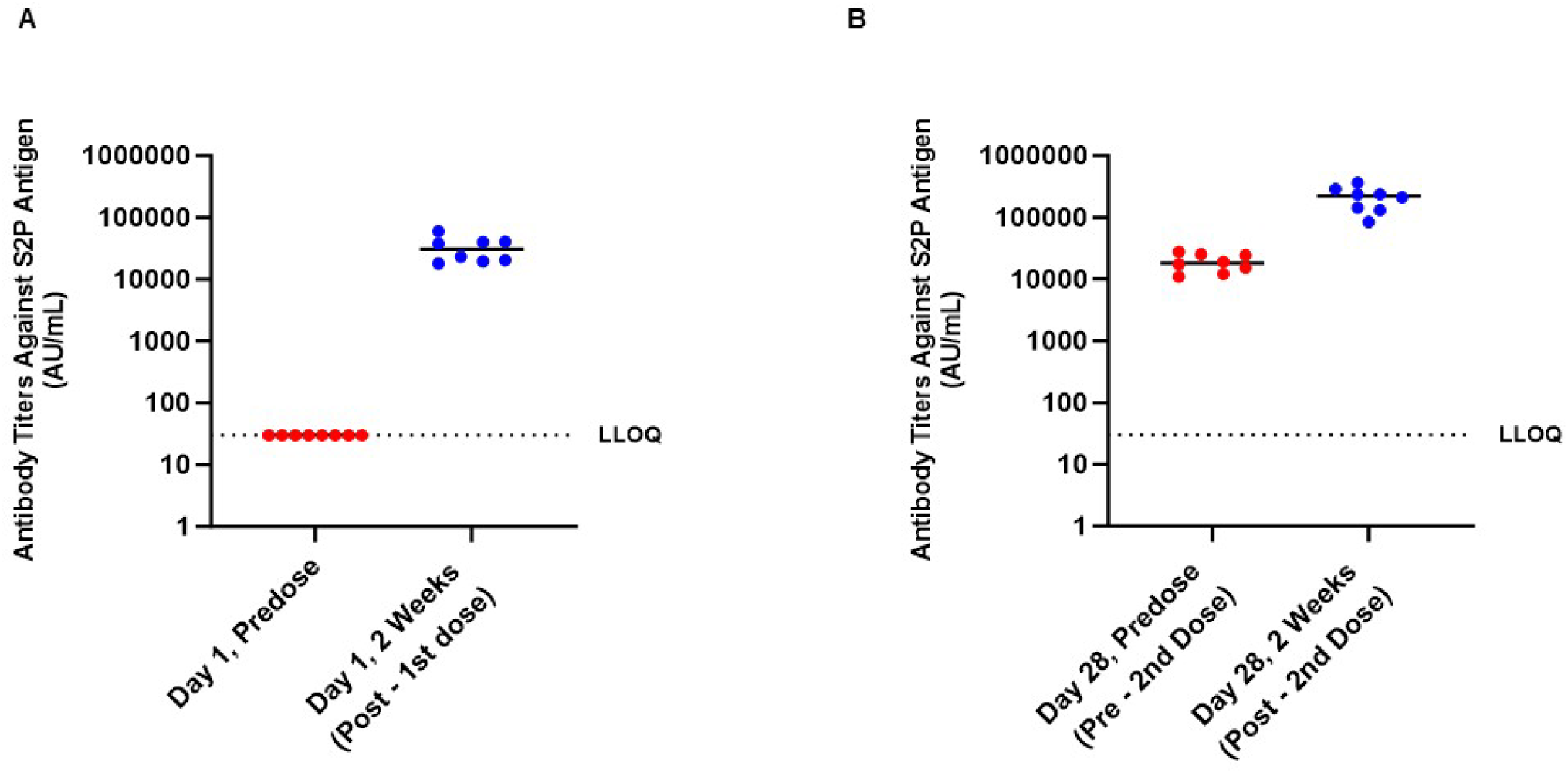
Antibody titers against SARS-CoV-2 S2P antigen following administration of mRNA-1273 in Sprague–Dawley rats. (A) Titers on Day 1 at pre-dose and 2 weeks post–first dose. (B) Titers on Day 28 at pre-dose (pre–second dose) and 2 weeks post–second dose. Individual animal values are shown with horizontal lines indicating group means. The dashed line represents the lower limit of quantification (LLOQ).

### Efficient lipid clearance via metabolism and excretion

Analysis of the Lipid H metabolic profile confirmed rapid clearance from systemic circulation and excretion through both hepatic and renal pathways. In plasma, parent Lipid H accounted for nearly all detectable signals at early timepoints, with only trace levels of metabolites present (Figure 4A). By contrast, metabolites dominated in urine and bile excreta (Figure 4B and 4C). In urine, multiple metabolites (mainly M1–M4) were observed within the first 24 h post-dose, with negligible amounts of parent Lipid H detected, indicating efficient renal elimination through biotransformation (Figure 4B). A fraction of parent Lipid H was observed in bile; however, metabolites (M1–M12) were the dominant species, representing the majority of the relative abundance at later intervals (Figure 4C). Only small amounts of parent Lipid H were detected in excreta, indicating that clearance occurred primarily through metabolism rather than direct elimination of the parent Lipid H.

**Figure 4.**
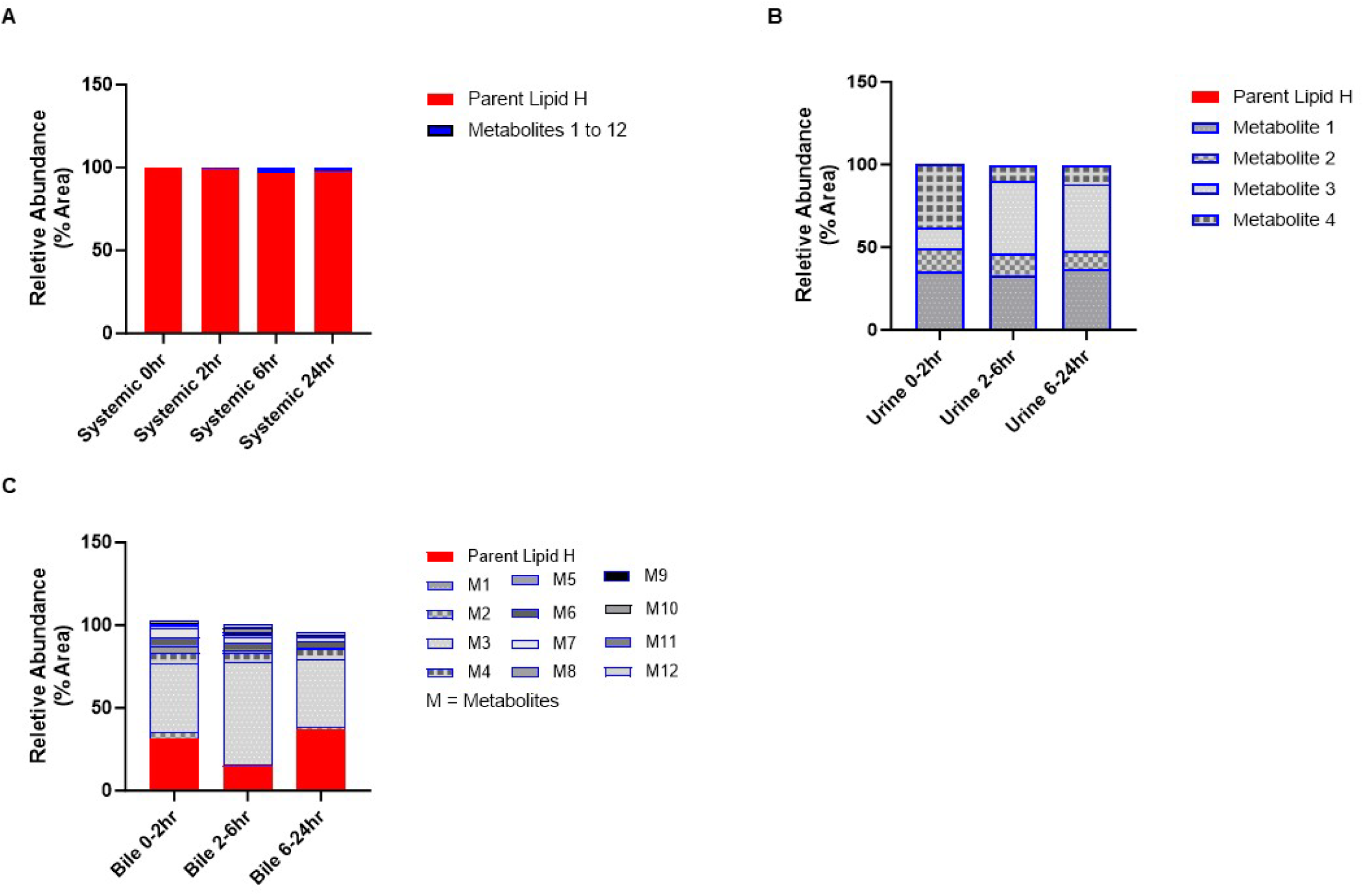
Relative abundance of parent lipid H and metabolites in Sprague–Dawley rats. (A) Systemic circulation at 0–24 h post-dose. (B) Urine collected over 0- to 2-h, 2- to 6-h, and 6- to 24-h intervals. (C) Bile collected over 0- to 2-h, 2- to 6-h, and 6- to 24-h intervals. Bars represent the relative abundance (% area) of parent Lipid H and identified metabolites (M1–M12). Metabolites were quantified by LC–MS/MS and expressed as percentage of total peak area.

Metabolite identification in rat plasma, urine, and bile identified 12 distinct metabolites derived from pathways including ester hydrolysis, β-oxidation, hydroxylation, and glutathione conjugation (Supplemental Table S2). *In vitro* studies using hepatocytes from human, rat, and nonhuman primates confirmed that these pathways were conserved across species, with consistent detection of major metabolites M1, M3, M4, M6, and M7; no human-specific metabolites were detected (Supplemental Table S3).

## Discussion

mRNA–LNP vaccines are now licensed worldwide for the prevention of several respiratory diseases, with a growing clinical footprint expanding into other indications. When considering Moderna’s approved mRNA-1273 (Spikevax™) and Pfizer–BioNTech’s BNT162b2 (Comirnaty™), acceptable safety and effectiveness profiles have been demonstrated and are attributed to the prevention of millions of hospitalizations and deaths due to Covid-19.^20–23^ This extensive real-world experience has established mRNA-LNP vaccine technology as a reliable and adaptable platform. Using rigorous methods that have been accepted in regulatory filings and adhere to international regulatory guidelines, here we report a comprehensive, cross-product dataset quantifying the biodistribution of mRNA-LNP vaccines. This analysis was conducted using 3 IM-administered mRNA-LNP drug products formulated in an identical LNP matrix (mRNA-1273, mRNA-1647, and a reporter mRNA [NPI-Luc] formulated in Lipid H LNPs), in Sprague Dawley rats at conservatively high (>130-fold human) dose levels to match toxicologic doses (when normalized to differences in body weight). Thus, the results represent a conservative and extreme estimate of tissue and systemic exposure of mRNA, LNP components, and/or SARS-CoV-2 Spike protein.

Across these Moderna products, the tissues with the highest exposure were consistently the injection site, draining lymph nodes, and spleen, with minimal distribution to other organs. This biodistribution profile is consistent with results used to support other licensed vaccine platforms following IM dosing in rodents and/or rabbits, including Pfizer–BioNTech’s BNT162b2 mRNA-LNP vaccine,^17^ as well as adjuvanted protein, and viral-vector vaccines.^24–27^ This shared biodistribution pattern across vaccine modalities supports a conserved mechanism of immune activation that facilitates vaccine uptake and antigen presentation primarily within local and lymphoid compartments after IM administration. The lymphoid tropism of mRNA-LNP vaccines also likely reflects the optimized nanoparticle size (60–120 nm), surface charge, and apolipoprotein-E–mediated interactions that promote retention within draining lymph nodes.^28–31^ A previous report in non-human primates also demonstrated a lymphoid-focused pattern following IM administration of an mRNA-LNP product formulated in the same LNP matrix as the products used here,^32^ indicating cross-species similarities and supporting the translational relevance of the data to higher-order species.

mRNA and LNP components cleared rapidly (within 48 h in serum; <14 days in lymphoid tissues) and first-order elimination modeling of these analytes revealed that concentrations declined to below quantifiable or very low to negligible levels (i.e., near complete elimination) by ∼2 weeks in the tissue with the longest T_1/2_ (spleen). Furthermore, there was no accumulation of mRNA or Lipid H in the systemic circulation or tissues following repeat dosing of mRNA-1273, providing additional evidence that there is no persistence of mRNA or Lipid H in either location.

When considering AUC, exposures of mRNA and Lipid H in non-lymphoid tissues were minimal, representing only a small fraction of that observed at the injection site. Albeit at low levels, the liver exhibited the greatest exposure among distal tissues (<3% relative to the injection site). In contrast, other tissues displayed only trace amounts compared with the injection site, which serves as the primary depot following IM administration. In support of liver tropism, published data using a different ionizable lipid (Lipid 5) via the intravenous (IV) route (which would yield much higher systemic exposure than IM), demonstrated similar rapid distribution almost exclusively to the liver and intestines by 24 h post-dose, supporting the expected clearance behavior of LNPs upon systemic entry.^33^

Across studies, the trace but detectable levels of mRNA or Lipid H in distal tissues, such as heart, lung, and brain, were near the assay’s LLOQ and/or were highly variable. Specifically, quantifiable levels in these tissues fluctuated across studies, sexes, timepoints, and even within use of the same drug product. Related to the last point, for example, following administration of mRNA-1647, concentrations of the 6 different mRNAs exhibited discordant levels across these low-concentration tissues. There were also occasional detections of mRNA without corresponding Lipid H, which are unlikely to represent true exposure since unprotected RNA is rapidly degraded in biological fluids and thus not expected to persist in tissues without LNP encapsulation.^32, 34^ These low-level signals are therefore attributed to analytical noise near the LLOQ or to residual vascular content after perfusion, rather than a physiologically relevant distribution. Perfusion substantially reduces intravascular content but cannot completely clear capillary beds; residual blood can thus account for trace detections near the assay limit. Additionally, trace brain mRNA signals are not considered indicative of blood-brain barrier penetration, given the restrictive barrier architecture, the LNP size (∼60–120 nm), and the lack of active transport ligands of LNPs.^35–38^

Discrepancies in tissue kinetics were occasionally noted between Lipid H and encapsulated mRNAs, which can be attributed to both biological and technical factors. First, the intracellular degradation of mRNA and lipids are different. Specifically, mRNA is inherently more labile than lipids, and is subject to nuclease degradation and endosomal processing,^39, 40^ whereas lipid clearance follows distinct esterase-mediated pathways. Second, differences in the LLOQ of the bioanalytical methods used to measure mRNA and lipid could also result in variable kinetics. Despite these differences, the tissues with the highest mRNA exposure were also always those with the highest Lipid H exposure. Thus, for mRNA-LNP drug products, these results provide scientific basis for using either analyte as a practical surrogate for the other when the goal is to compare tissue exposure across organs.

To our knowledge, this is the first characterization of SARS-CoV-2 Spike protein exposure and clearance in a SARS-CoV-2–naïve model, addressing a key limitation of human studies where pre-existing or subclinical SARS-CoV-2 infection cannot be excluded. We found that in rats, where there were no prior existing antibodies against SARS-CoV-2 prior to administration mRNA-1273, Spike protein is transient in the systemic circulation. This is demonstrated by a serum T_1/2_ of ∼18 h (albeit calculable only after a first dose), >98% simulated elimination within 5 days post-dose, and lack of accumulation following a second dose of mRNA-1273. Recent claims that Spike protein persists for months following vaccination are either not aligned with the broader body of scientific evidence or have technical limitations that may confound interpretation. For example, Bhattacharjee et al.^41^ relied on small, uncontrolled samples, and most importantly, used bioanalytical methods that could not distinguish vaccine-derived Spike protein from Spike protein from prior or ongoing COVID-19 infection. Additionally, Patel et al.^42^ asserted persistence of Spike protein in non-classical CD16+ monocytes for more than 200 days post-vaccination, which is inconsistent with the recognized ∼7-day lifespan of these cells. A more plausible explanation for these latter findings is that intermittent detection of Spike protein reflects latent SARS-CoV-2 infection and/or ongoing viral infection.^43^ Altogether, our data, derived from a well-controlled animal study, along with a clinical study,^44^ do not support persistence of vaccine-derived Spike protein beyond 5 days after vaccination.

AUEC_last_ for systemic Spike protein levels was estimable after the first dose, but not after the second dose, due to insufficient quantifiable serum timepoints. The most plausible explanation for this result is related to the bioanalytical method used to detect Spike protein in serum rather than a true lack of protein expression following repeat dosing. Specifically, the enzyme-linked immunosorbent assay (ELISA) method used for Spike protein quantification detected only unbound (free) antigen; therefore, the robust, SARS-CoV-2 S2P-specific polyclonal antibodies elicited after the first dose likely bound to newly expressed Spike protein, forming immune complexes that masked epitopes recognized by the capture and detection antibodies. This potential interference could have contributed to the apparent lack of detectable Spike protein after the booster dose and warrants further investigation using orthogonal detection methods (e.g., liquid chromatography-mass spectrometry [MS]).

Lipid H underwent rapid and predictable biotransformation with intact lipid representing the predominating circulating species after IV dosing in Sprague Dawley rats. Ester hydrolysis and β oxidation were the primary metabolic pathways observed *in vivo* and *in vitro*, generating metabolites that were efficiently cleared through renal and biliary routes. Low-molecular-weight, hydrophilic metabolites were especially enriched in urine, consistent with renal filtration. Taken together, the extensive metabolism of Lipid H, rapid overall clearance (within 24 h) and the efficient elimination of both the parent and its metabolites to <3% of the maximum level indicate that Lipid H is unlikely to accumulate upon repeat IM dosing, even in the context of patients with hepatic or renal impairment.

Using Sprague Dawley rats as the model system enabled us to comprehensively characterize the mRNA, Lipid H, and expressed antigen exposure time courses in both blood and tissues, the latter of which are measurements that are not feasible to evaluate in humans. However, there are important limitations that should be acknowledged. Rats are not humans, and the doses that were used were supratherapeutic (>130-fold human when normalized to species body weight). While such dosing and species differences may influence quantitative pharmacokinetics (PK) and likely overestimate systemic and tissue exposure relative to humans, qualitative biodistribution and elimination patterns are expected to be generally conserved across species. Recently published human PK data for Lipid H in systemic circulation following mRNA-1273 vaccination help support this conclusion. In that study, Kent et al.^45^ reported that Lipid H is cleared from systemic circulation within 2 weeks post-vaccination. This longer human T_1/2_ (∼27 h) relative to what was observed in rats based on our data (∼6 h) is not unexpected because rodents generally metabolize drugs more rapidly than humans. Another important consideration is that the last blood and tissue collections across studies occurred up to 14 days post-dose. Consequently, modeling was employed to estimate the later phase of elimination, a standard approach for characterizing decay kinetics when data are truncated.^46^ The model’s predictions matched the overall patterns seen with reported human data (which was evaluated out to 4-weeks after vaccination)^45^, suggesting that the rat results remain reliable even with this limitation.

In conclusion, by harmonizing the dose level, route of administration, and LNP composition, and incorporating an evaluation of distribution following repeat dosing, we provide a comprehensive view of the metabolism, clearance, and tissue distribution of mRNA-LNP vaccines. The encoded Spike protein or immune response did not alter tissue tropism, and exposures for all vaccine components were transient with no evidence of persistence beyond 2 weeks. Furthermore, the metabolism of Lipid H favored clearance over storage. The consistent distribution and clearance patterns across products provide a weight of evidence to justify bridging biodistribution data when products share the same LNP composition and route of administration. This approach aligns with the risk-based framework endorsed by major regulatory agencies (e.g., European Medicines Agency, United States Food and Drug Administration [FDA], Japan’s Pharmaceuticals and Medical Devices Agency, and the World Health Organization) and supports platform-level, read-across for LNP-formulated nucleic acid medicines. As such, these findings reinforce ongoing efforts to reduce redundant *in vivo* testing consistent with the 3 Rs (replacement, reduction, and refinement), and the FDA’s roadmap promoting scientifically justified non-animal strategies to support drug development.^47^

## Materials and Methods

### Animals and ethical compliance

All animal care and experimental procedures were conducted in compliance with the Guide for the Care and Use of Laboratory Animals and were approved by the Institutional Animal Care and Use Committee (Charles River Laboratories, Inc, Mattawan, MI, USA or Worcester, MA, USA) or by the ethical committee of Charles River Laboratories Safety Assessment Services.

### mRNA-LNP products

Biodistribution studies were conducted using three mRNA–LNP products, where mRNA was encapsulated in the same 4 lipids (Lipid H, PEG2000-DMG, cholesterol, DSPC) with generally similar ratios of lipids to mRNA: (1) mRNA-1273 (Spikevax™), Moderna’s licensed SARS-CoV-2 vaccine; (2) mRNA-1647, an investigational CMV vaccine candidate (containing 6 mRNA constructs); and (3) an mRNA–LNP product that encodes a reporter protein, NPI-Luc. For the metabolism and excretion studies, a product encapsulating a non-translating mRNA was used (non-translating Factor IX or NT-FIX), which was formulated similarly to the other products in the same 4-lipid matrix. All dose levels listed in the *in vivo* studies are reflective of the RNA dose.

The test materials were supplied preformulated. While the study with mRNA-1273 and NPI-Luc used the formulations as received, the study with mRNA-1647 used phosphate-buffered saline (PBS) dilution to achieve target dose concentration.

Of the 4 lipids in the LNP formulation, Lipid H was the only one evaluated in the biodistribution, metabolism, and excretion studies for the following reasons: (1) cholesterol is widely present throughout the body and therefore it is not possible to differentiate drug product-delivered cholesterol exposure from endogenous, naturally occurring cholesterol, and (2) DSPC and PEGylated molecules are lipids already in approved drug products^48^ at doses in excess of those in the clinical IM doses of the vaccines and/or as IV administered products (which would result in significantly greater systemic exposure than with IM administration).

### Experimental designs

The biodistribution studies evaluating mRNA-1273 and NPI-Luc in Lipid H-containing LNPs were conducted by Charles River Laboratories, Inc., Mattawan, MI, USA, and used male and female Crl:CD(SD) Sprague Dawley rats (Charles River Laboratories, Inc). Prior to initiation of dosing, the average age and weight range of animals in the mRNA-1273 study were 10.5–13.5 weeks and 225 g (females) and 408 g (males), respectively. In the NPI-Luc study, these biometrics traits ranged from 10 to 11 weeks of age and weights of 223 g (females) and 351 g (males). The biodistribution study evaluating mRNA-1647 was conducted by Charles River Laboratories Montreal ULC, Sherbrooke, QC, Canada, and used male Crl:CD(SD) Sprague Dawley rats (Charles River Laboratories, Inc). Prior to initiation of dosing, the age and average weight of animals in the mRNA-1647 study was 8 weeks and 323 g, respectively. The *in vivo* metabolism and excretion study with NT-FIX mRNA in Lipid H LNPs was conducted by Charles River Laboratories, Inc, Worcester, MA, USA, and used male Sprague Dawley rats (Envigo EMS, average weight 259 g at initiation of dosing).

In all the studies, animals were acclimated to the facilities for at least 6–7 days and randomly assigned into groups. Rats were housed in microisolator cages under controlled environmental conditions (12-h light/dark cycle, 18–26°C, 40–70% humidity) with *ad libitum* access to food (MI Nutrition International Certified Rodent Chow No. 5CR4, 14% protein or Teklad Global Diets™, 18% Protein Rodent Diet 2018) and locally sourced municipal tap water that was passed through reverse osmosis, ultraviolet irradiated, and provided *ad libitum*.

### In vivo biodistribution studies

The route of administration used to assess biodistribution was IM, consistent with the clinical route. All doses were administered via IM injection into the lateral thigh (quadriceps) in a 200-µL volume using a 27-gauge needle. In line with biodistribution regulatory guidance,^49^ the selected dose levels of the mRNA-LNP products bridged dose levels used in toxicology studies or were the maximum feasible dose based on the drug concentration and the maximum allowable dosing volume for IM administration (200 µL/dose).^49^

Table 7 summarizes the administered rat dose levels on both a flat per dose basis (e.g. µg/dose) and body-weight normalized basis (µg/kg) for each of the drug products as it provides the corresponding “safety margin” relative to clinical dose when normalized to the average body weight of each species. Once in circulation, interspecies disposition and PK of extravascularly administered drugs are governed primarily by body-size-related clearance.^50–52^ Therefore, given that these IM biodistribution studies are aimed at assessing exposure and kinetics systemically, it is most physiologically relevant to normalize dose based on body weight (µg/kg) when making rat–human comparisons. Thus, the studies intentionally exceeded clinical dose levels on µg/kg basis to enable comprehensive systemic biodistribution assessments at exaggerated exposures, in alignment with International Council for Harmonisation of Technical Requirements for Pharmaceuticals for Human Use S12 guidance.

**Table 7.**
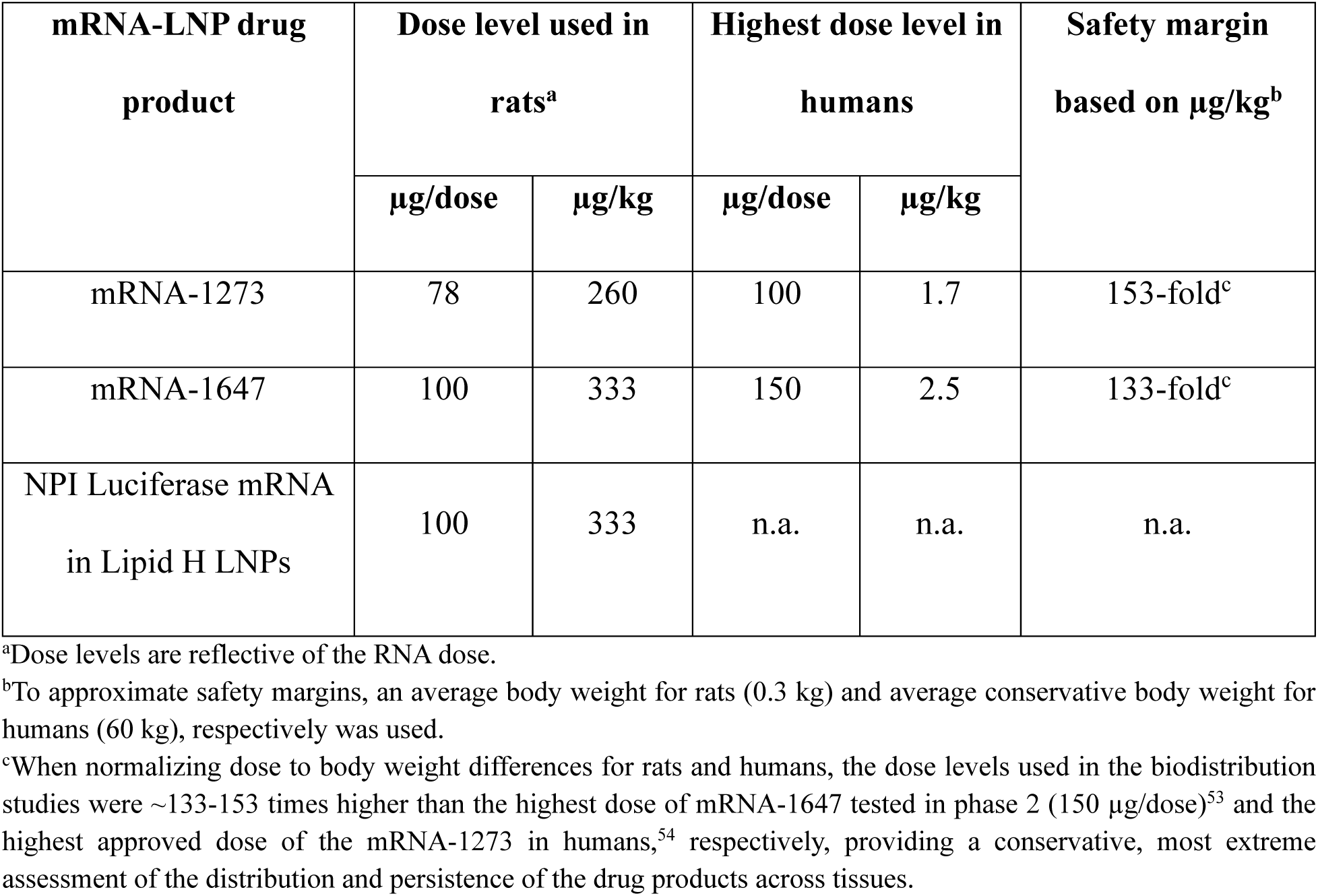
Dose levels used in rat biodistribution studies are >130-fold higher than human doses, representing a conservative, most extreme assessment of tissue distribution and exposure.

| mRNA-LNP drug<br>product | Dose level used in<br>rats<sup>a</sup> |  | Highest dose level in<br>humans |  | Safety margin<br>based on µg/kg<sup>b</sup> |
| --- | --- | --- | --- | --- | --- |
|  | µg/dose | µg/kg | µg/dose | µg/kg |  |
| mRNA-1273 | 78 | 260 | 100 | 1.7 | 153-fold<sup>c</sup> |
| mRNA-1647 | 100 | 333 | 150 | 2.5 | 133-fold<sup>c</sup> |
| NPI Luciferase mRNA<br>in Lipid H LNPs | 100 | 333 | n.a. | n.a. | n.a. |

<sup>a</sup>Dose levels are reflective of the RNA dose.
<sup>b</sup>To approximate safety margins, an average body weight for rats (0.3 kg) and average conservative body weight for humans (60 kg), respectively was used.
<sup>c</sup>When normalizing dose to body weight differences for rats and humans, the dose levels used in the biodistribution studies were ~133-153 times higher than the highest dose of mRNA-1647 tested in phase 2 (150 µg/dose)<sup>53</sup> and the highest approved dose of the mRNA-1273 in humans,<sup>54</sup> respectively, providing a conservative, most extreme assessment of the distribution and persistence of the drug products across tissues.

mRNA-1647 and NPI-Luc mRNA in Lipid H-containing LNPs were administered as a single IM dose. mRNA-1273 was administered IM twice, once on Day 1 and once contralaterally 4 weeks later (i.e., on Day 28) to evaluate tissue distribution after repeat administration and when using the most compressed clinical dosing regimen.

Four animals/sex/timepoint/dosing occasion were administered mRNA-1273, and 3 and 5 animals/sex/timepoint were administered NPI-Luc in Lipid H-containing LNPs and mRNA-1647, respectively. Animals were monitored daily for morbidity, mortality, clinical signs, and local injection site reactions. Body weights were recorded at baseline and prior to necropsy. In the study with mRNA-1273, body weights were also recorded weekly recordings post dose of mRNA-1273 No unscheduled deaths occurred, and all animals completed the in-life phase as planned.

### Tissue collection and processing

Following administration of 1 or 2 doses of mRNA-1273, whole blood and tissues were collected at 0.16, 1, 4, 10, 24, 72, 120, 168, and 336 h post-dose. Following administration of NPI-Luc mRNA in Lipid H-containing LNPs, whole blood and tissues were collected at 0.16, 1, 4, 10, 24, 72, 120, and 168 h post-dose. Following administration of mRNA-1647, whole blood and tissues were collected at 0 (pre-dose), 2, 8, 24, 48, 72, and 120 h post-dose for mRNA-1647 quantitation.

Animals were euthanized by deep isoflurane inhalation at predefined time points and were perfused with sodium chloride 0.9%, heparin (1000 IU/mL), 1% sodium nitrite, and PBS until the fluid runs were sufficiently clear. Tissues were then excised rapidly, rinsed in cold PBS, blotted dry, weighed, and snap-frozen in liquid nitrogen. Whole blood was collected via cardiac puncture into K₂-EDTA tubes for plasma or serum separation.

Collected tissues across the three studies primarily included: bone marrow, brain, eye, heart, injection site muscle, jejunum, kidney, liver, lymph nodes (axillary, inguinal, and popliteal), ovary, spleen, stomach, testes, thymus, and uterus. The selected tissues reflect both regulatory expectations for the IM route of administration and data derived previously from the developer demonstrating exposure (or lack thereof) in relevant tissues. Differences in tissues collected in the studies are noted in the Results section in the appropriate tables.

### Bioanalytical methods for biodistribution studies

A summary of the bioanalytical methods and assay rigor used for the biodistribution experiments is provided in Table 8 with detailed methods described below.

**Table 8.** Analytical methods and assay rigor used for rat biodistribution studies, in vivo metabolism and excretion study, and in vitro metabolism study.

| Analyte | Analytical Method | Species | Matrix | Assay Rigor | Calibration Range |
| --- | --- | --- | --- | --- | --- |
| NPI-Luc mRNA | bDNA Single plex | Rats | Serum, tissues | Qualified | 0.125 to 8.00 ng/mL |
| mRNA from mRNA1273 | RT-qPCR | Rats | Serum, tissues | Qualified | 0.850 to 170,000 pg/mL for serum<br>0.850 to 170,000 pg/g for tissue |
| mRNAs from mRNA1647 (6 different mRNAs) | bDNA multiplex | Rats | Plasma, tissues | Fit for purpose | 0.01 to 5 ng/mL for 4 mRNAs<sup>a</sup><br>0.05 to 5 ng/mL for 2 mRNAs<sup>a</sup> |
| Lipid H | LC-MS/MS | Rats | Plasma, tissues | Qualified | 0.500 to 500 ng/mL |
|  |  |  | Plasma, bile, urine | Fit for purpose | 0.2 to 500 ng/mL |
| NPI-Luc protein | ECL | Rats | Tissue | Qualified | 200 to 16,000 ng/g |
| SARSCoV2 S protein | LBA | Rats | Serum | Validated | 4.57 to 10,000 ng/mL |
|  |  |  | Tissues | Qualified | 91.4 to 100,000 ng/mL |
| Anti-SARS-CoV-2 S protein IgG antibody | ELISA | Rats | Serum | Fit for purpose | 0.059 to 186.3 Antibody Unit/mL<sup>b</sup> |
| Lipid H metabolite identification | LC-HRMS | Rat | Plasma, bile, urine | Fit for purpose | NA |
| Lipid H metabolite identification | LC-HRMS | *In vitro* | Cryopreserved hepatocytes from Sprague Dawley rats, cynomolgus monkey, and human | Fit for purpose | NA |

Abbreviations: AU = arbitrary units; bDNA = branched DNA; ECL = electrochemiluminescence ligand binding assay; ELISA = enzyme-linked immunosorbent assay; ESI = electrospray ionization; HRMS = high-resolution mass spectrometry; IgG = immunoglobulin G; LBA = ligand binding assay; LC = liquid chromatography; LLOQ = lower limit of quantification; Luc = luciferase; mRNA = messenger RNA; MS = mass spectrometry; MS/MS = tandem mass spectrometry; NA = not available; NPI = nascent peptide imaging; PEG2000-DMG = 1,2-dimyristoyl-rac-glycero-3-methylpolyoxyethylene glycol-2000; RT-qPCR = quantitative reverse transcription polymerase chain reaction; SARS-CoV2 = severe acute respiratory syndrome coronavirus;
Note: For the calculation of tissue-to-plasma/serum ratios, where tissue concentration is in ng/g and plasma/serum concentration is in ng/mL, the assumption is that 1 g of tissue is equivalent to 1 mL of plasma/serum.
<sup>a</sup> LLOQ reflects the standard range in neat plasma and tissue homogenate solution (after applied 100x minimum required dilution)
<sup>b</sup>The analytical lower limit of quantification (LLOQ) was 0.059 Antibody Unit (AU)/mL Samples were assayed at a minimum required dilution (MRD) of 1:500, resulting in a reportable LLOQ of 30 AU/mL (0.059 × 500) in undiluted serum.

### mRNA quantification

Reverse transcription-quantitative polymerase chain reaction (RT-qPCR) was used for quantification in the mRNA-1273 study and reflects the most sensitive analytical method for detecting RNA in biological matrices. Single-plex branched DNA (bDNA) hybridization and multiplex bDNA bead-based detection pre-dated use of RT-qPCR and were used for the NPI-Luc in Lipid H-containing LNP and mRNA-1647 studies, respectively. Despite these differences in methodology, each method was qualified for precision, accuracy, selectivity, and sensitivity.

Quantification of the mRNA in mRNA-1273 was performed using a one-step RT-qPCR assay. Tissues were homogenized prior to RNA isolation in Maxwell RSC simplyRNA homogenization buffer. Total RNA was extracted from tissue homogenates and serum using the Maxwell RSC simplyRNA isolation procedure. Complementary DNA was synthesized using reverse transcriptase and amplified with TaqMan probe sets specific to the mRNA-1273 sequence on a real-time polymerase chain reaction system. Calibration curves were generated using known concentrations of mRNA-1273 drug product with curve range from 0.00085 to 170 ng/mL in rat serum and 0.00085 to 170 ng/g in tissue.

Levels of mRNA from NPI-Luc in Lipid H-containing LNPs were quantified using a single-plex bDNA hybridization assay (QuantiGene® 2.0 assay system, Thermo Fisher Scientific). Tissue homogenates and serum samples were lysed in the presence of proteinase K and hybridized with sequence-specific capture, label, and blocker probes targeting the luciferase transcript. Chemiluminescent signals were generated following amplification with alkaline phosphatase–conjugated probes and substrate and measured on a SpectraMax M3 luminometer. The calibration curve ranged from 0.125 to 8.00 ng/mL in rat serum. mRNA levels in each rat tissue homogenate were calculated against a single calibration curve prepared from pooled tissue homogenates with a calibration range of 0.125 to 16.0 ng/g. The pooled homogenates were generated by combining equal proportions of rat liver, spleen, heart, lung, kidney, brain, uterus, ovary, testes, eye, jejunum, axillary lymph node, tail, skeletal muscle, stomach, and thymus. The levels of the 6 different mRNA in mRNA-1647 were quantified using a multiplex bDNA hybridization assay (QuantiGene® Plex, Thermo Fisher Scientific/Affymetrix) run on a Luminex Bio-Plex Suspension Array system. This assay simultaneously quantified all 6 mRNAs (gB, gH, gL, UL128, UL130, and UL131A). Plasma and perfused tissue samples were lysed, homogenized, and hybridized with construct-specific capture, label, and blocker probes conjugated to distinct bead regions. Chemiluminescent signals were amplified through pre-amplifier, amplifier, and streptavidin-phycoerythrin–labeling steps, then detected bead-by-bead on the Luminex platform. The multiplex design allowed assessment of each construct in parallel and demonstrated highly concordant PK across constructs. The calibration curve ranged from 0.01 to 5 ng/mL in plasma and tissue homogenates, depending on the construct.

### Lipid H quantification

Quantification of Lipid H in plasma and tissue samples was conducted using a qualified liquid chromatography-tandem MS method. Tissues were homogenized in PBS and subjected to protein precipitation using a 1:1 mixture of acetonitrile and methanol. After centrifugation, the supernatant was transferred to 96-well plates for analysis.

Chromatographic separation was achieved on a C8 reversed-phase column (50 mm×2.1 mm, 5-µm particle size) using gradient elution with mobile phases consisting of aqueous ammonium formate and acetonitrile. A 10-µL injection volume was used, and each run had a total time of 4 min. The detection was performed using a Sciex API 5000 mass spectrometer equipped with electrospray ionization in positive ion mode.

Calibration curves were generated in matrix-matched standards and covered a concentration range of 0.5–500 ng/mL (or ng/g), with linear regression and appropriate weighting.

### Encoded protein quantification

Levels of the encoded SARS-CoV-2 Spike protein and NPI-Luc protein were quantified using a qualified ELISA method. For the SARS-CoV-2 Spike protein assay, the ELISA method was designed to measure the concentration of free Spike protein present in the serum and tissue samples.

Quantification of expressed NPI-Luc protein was performed using qualified electrochemiluminescence-based ligand-binding assays. Tissue homogenates were prepared and diluted to a minimum required dilution (MRD) in assay buffer before analysis. Plates were coated with hamster anti-V5 antibody, followed by sequential incubation of the sample with anti-firefly luciferase and SulfoTag-labeled secondary antibodies to generate an electrochemiluminescent signal proportional to analyte concentration. The assay demonstrated acceptable accuracy, precision, and selectivity, with a LLOQ of 200 ng/g in tissue homogenates.

Quantification of the expressed SARS-CoV-2 Spike protein following mRNA-1273 administration was performed using qualified colorimetric ligand-binding (sandwich ELISA) assays. The assays employed mouse anti-SARS-CoV-2 whole Spike protein as the capture antibody and biotin-conjugated rabbit anti-Spike antibody for detection, followed by streptavidin–peroxidase for signal generation. Serum samples were analyzed following an MRD of 1:20, while tissue homogenates were analyzed following an MRD of 1:10. The assays demonstrated acceptable accuracy, precision, and selectivity, with LLOQ of 4.57 ng/mL in serum and 91.4 ng/g in tissue homogenates.

### Antibody titers against SARS-CoV-2

The immunogenicity of mRNA-1273 was evaluated by assessing binding antibody titers against SARS-CoV-2. High-binding 96-well plates (MaxiSorp) were coated overnight (2–8°C) with SARS-CoV-2 S2P protein (GenScript; 1.5 µg/mL in Dulbecco’s PBS), washed 3× with Dulbecco’s PBS–0.05% Tween-20, and blocked with StartingBlock (PBS) for ≥45 min at room temperature. Rat serum standards and samples, diluted in StartingBlock T20 (PBS), were added (100 µL/well) and incubated for 2 h. After washing, goat anti-rat immunoglobulin(H+L)–horseradish peroxidase (KPL/SeraCare; 1:7,000 in StartingBlock T20) was applied for 1 h, plates were washed, and 3,3’,5,5’-tetramethylbenzidine (SureBlue Reserve) was developed for 30 min protected from light. Absorbance was read at 650 nm (VersaMax), and antibody units (AU)/mL were calculated by 4-parameter logistic interpolation from a standard curve generated using serial dilutions (1:102 to 1:10⁷) of a pooled hyperimmune rat serum reference standard. The analytical LLOQ was 0.059 AU/mL, corresponding to the lowest dilution of the standard producing signal ≥3.5-fold above background (≈1:3.2 × 10⁵ dilution). Samples were tested at an MRD of 1:500. Values below the analytical LLOQ (0.059 AU/mL) correspond to a reportable LLOQ of 30 AU/mL in undiluted serum. Samples exceeding the upper calibration standard were further diluted as necessary to fall within the validated range, consistent with standard ELISA practice.

### Metabolism and excretion studies

A summary of the bioanalytical methods and assay rigor used for the metabolism and excretion experiments is provided in Table 8 with detailed methods described below.

### In vitro metabolite identification and profiling of Lipid H

Primary rat, monkey, and human hepatocytes were thawed and resuspended in Williams’ E medium at 1×10^6^ cells/mL, with 250 µL of cell suspension or media used per condition; heat-denatured controls were prepared by boiling hepatocytes at 100 °C for 15 min. A Lipid H LNP stock solution was diluted in media to 100 µM and further diluted in cell suspensions to achieve a final concentration of 10 µM. Samples were collected at 0, 4, and 24 h, and Lipid H and its metabolites were extracted with 250 µL cold acetonitrile, centrifuged at 10,000 × *g* for 10 min at 10 °C, and the supernatant was removed for analysis. Metabolite profiling was performed on an Agilent 1290 UPLC coupled to an Agilent 6550 QTOF mass spectrometer using a reverse-phase C18 column, with data acquired in positive ion mode and instrument parameters optimized with a Lipid H standard; both data-dependent and targeted tandem MS were used for metabolite identification.

### In vivo metabolism and excretion study

The IV route of administration was used in the *in vivo* study to maximize systemic exposure and ensure the ability to detect Lipid H parent and metabolites in plasma, bile, and urine. Route of administration would not be expected to influence metabolism or elimination routes; thus, the data derived using IV dosing is also considered supportive of IM dosing. One group of male Sprague Dawley rats (n=3; bilateral jugular-vein cannulated/bile duct cannulated) was administered NT-FIX mRNA in Lipid H LNPs at a dose of 0.672 mg/kg (dose concentration 0.48 mg/mL; dose volume 1.4 mL/kg, diluted in buffer with Tris/Sucrose/Acetate, pH7.5) via a 10-min infusion followed by a 0.5 mL post-dose flush. Serial blood samples were collected from each animal prior to dosing and at 2, 6, and 24 h post-infusion start. Blood samples, which were collected into tubes containing K_2_-EDTA, were processed for plasma. Urine and bile were collected from 0 to 2 h, 2 to 6 h, and 6 to 24 h post-infusion start. Metabolite profiling was conducted by liquid chromatography–high-resolution MS.

### Kinetic analysis, reporting, and simulation of concentration–time profile

#### PK and PD evaluation

PK and PD parameters were estimated using noncompartmental analysis in Phoenix PK software. For PK assessments (mRNA and Lipid H), concentration–time data in plasma or serum, respectively, and all tissues except the injection site were analyzed using an extravascular model. The IV bolus model was applied for the injection site. PK parameters were appropriately calculated based on either lipid or mRNA dose. For the kinetic evaluation of mRNA-1273, all parameters were generated from composite Lipid H, and mRNA concentrations collected on Days 1 and 28, whenever available, assuming a pre-dose concentration of zero on Day 28 for the extravascular route of administration.

For the PD assessment (expressed protein), concentration–time profiles of the encoded protein (NPI-Luc or SARS-CoV-2 Spike protein) in serum and tissues were analyzed using the same non-compartmental approach consistent with the extravascular route of administration. For the study involving mRNA-1273, the Day 28 pre-dose concentration was also assumed to be zero since the previous collection concentration was below the LLOQ.

For both PK and PD analyses, concentrations below the LLOQ were treated as zero, and concentrations above the limit of quantitation were treated as the upper limit of quantitation (ULOQ) for the purposes of the PK data analysis. Comprehensive estimation of primary or secondary PK parameters (including C_max_, time to maximum concentration [T_max_], area under the concentration-time curve from time 0 to time t [AUC_(0-t)_], area under the concentration-time curve to the last measurable concentration [AUC_last_], area under the concentration-time curve from time 0 to infinity [AUC_(0-inf)_], time of the last measurable concentration [T_last_], effective T_1/2_, mean residence time to the last measurable concentration [MRT_last_], clearance normalized by bioavailability [Cl/F], apparent volume of distribution during the terminal phase after non-IV administration [V_z_/F], tissue to plasma/serum ratio (%) of AUC, tissue to plasma/serum ratio (%) of C_max_), and PD parameters (including E_max_, TE_max_, area under the effect-time curve from time 0 to time t [AUEC_(0-t)_], AUEC_last_, time of the last measurable effect [TE_last_], effective T_1/2_, MRT_last_) were calculated for each of the studies. While all PK and PD parameters were calculated in the studies to support regulatory submissions, the following parameters were selected for reporting in this manuscript as they best characterize tissue exposure and relative persistence and clearance: AUC_(0-t)_, AUC_last_, tissue to plasma/serum ratio (%) of AUC and effective T_1/2_ for PK, and E_max_, AUEC_last_, TE_last_ and effective T_1/2_ for PD.

Tissue-to-systemic exposure ratios (%) were calculated as (AUC_tissue_/AUC_systemic_) × 100. AUC was defined as the area under the concentration versus time curve from the start of dose administration to the last observed quantifiable concentration using the linear trapezoidal method to calculate.

### Combined sex kinetic data reporting

To characterize broad-stroke kinetic estimates with the mRNA-1273 and NPI-Luc mRNA in Lipid H-containing LNPs dataset, we pooled male and female PK and PD data (otherwise referred to as “Sex Combined”), except where tissues were sex-dependent (e.g., testes or ovaries). Combining male and female data increased sample size, effective power and avoided over-partitioning the dataset. Sex differences were, however, always evaluated as a potential covariate in the studies. There were no overall sex differences in individual serum, plasma, and or tissue mRNA concentrations in the study with NPI-Luc mRNA in Lipid H-containing LNPs (Supplemental Table S1), which was only administered as a single dose. In the study with mRNA-1273, where two dose administrations were administered over the course of 1 month and significant sex-dependent weight difference existed, mRNA exposures in some matrices were higher in females than males (Supplemental Table S1). The differences in exposure are attributed to differences in body weight: female body weights were approximately 1.5- to 2-fold lower than males across dosing days (Supplemental Figure S1). As a result, females received proportionally 1.5- to 2-fold higher doses of mRNA-1273 per kg of body weight than males, providing the most likely explanation for the sex-related differences in exposure.

### Simulation of mRNA and Spike protein elimination profiles

Simulated concentration–time profiles for mRNA and spike protein were generated using observed C_max_ or E_max_ values and tissue- or plasma-specific effective T_1/2_. First-order elimination kinetics were assumed, with concentrations estimated using Ct = C_max_ or E_max_×e^−kel*t^ and k_el_ = ln (2)/t_1/2_. Profiles were simulated over six half-lives to illustrate the extent of elimination, corresponding to ∼1.6% of C_max_ or E_max_ remaining (98.4% eliminated). Simulations and curve fitting were conducted in GraphPad Prism (GraphPad Software, San Diego, CA).

### Statistical analysis

To assess differences in exposure between the first and second dose of mRNA-1273, data are presented as mean values ± standard deviation (SD) from two independent dosing days (Figure 2). Comparisons between dosing days were performed using an unpaired Student’s *t*-test. Statistical analyses were conducted using GraphPad Prism (GraphPad Software, San Diego, CA, USA). A *p* value of less than 0.05 was considered statistically significant.

## Supporting information

Goody_Supplement

## Data Availability Statement

Supporting data may be made available from the corresponding author upon reasonable request

## Acknowledgements

John Wickwire, Harkewal Singh, Saara Mansouri, Matthew Gorman, Shyam Kumar Gudey, Kelin Wang, and Charles River Laboratories. Medical writing and editorial assistance were provided by MEDiSTRAVA in accordance with Good Publication Practice guidelines, funded by Moderna, Inc., and under the direction of the authors.

## Author Contributions

SMGG, CR, YL, and NC contributed to the study concept and design. SMGG, CR, YL, and NC analyzed and interpreted the data. All authors contributed to the drafting and critical review of this manuscript and approved the final draft.

## Declaration of Generative AI and AI-assisted technologies in the writing process

During manuscript drafting, the authors used ChatGPT for language editing and phrasing suggestions only. No AI tools were used for data analysis, interpretation, or generation of scientific conclusions, and all final text was reviewed and approved by the authors.

## References

1. Hassett, K. J., Benenato, K. E., Jacquinet, E., Lee, A., Woods, A., Yuzhakov, O., Himansu, S., Deterling, J., Geilich, B. M., Ketova, T., et al. (2019). Optimization of lipid nanoparticles for intramuscular administration of mRNA vaccines. Mol. Ther. Nucleic Acids 15: 1–11. 10.1016/j.omtn.2019.01.013.

2. Gebre, M. S., Brito, L. A., Tostanoski, L. H., Edwards, D. K., Carfi, A., and Barouch, D. H. (2021). Novel approaches for vaccine development. Cell 184: 1589–1603. 10.1016/j.cell.2021.02.030.

3. Pardi, N., Hogan, M. J., Porter, F. W., and Weissman, D. (2018). mRNA vaccines—a new era in vaccinology. Nat. Rev. Drug Discov. 17: 261–279. 10.1038/nrd.2017.243.

4. Espeseth, A. S., Cejas, P. J., Citron, M. P., Wang, D., DiStefano, D. J., Callahan, C., Donnell, G. O., Galli, J. D., Swoyer, R., Touch, S., et al. (2020). Modified mRNA/lipid nanoparticle-based vaccines expressing respiratory syncytial virus F protein variants are immunogenic and protective in rodent models of RSV infection. NPJ Vaccines 5: 16. 10.1038/s41541-020-0163-z.

5. Victora, G. D., and Nussenzweig, M. C. (2022). Germinal centers. Annu. Rev. Immunol. 40: 413–442. 10.1146/annurev-immunol-120419-022408.

6. US Food and Drug Administration (2022). Summary Basis for Regulatory Action - SPIKEVAX https://www.fda.gov/media/155931/download.

7. US Food and Drug Administration (2024). Summary Basis for Regulatory Action - MRESVIA https://www.fda.gov/media/179634/download?attachment.

8. US Food and Drug Administration (2021). Summary Basis for Regulatory Action - COMIRNATY https://www.fda.gov/media/151733/download.

9. Soens, M., Ananworanich, J., Hicks, B., Lucas, K. J., Cardona, J., Sher, L., Livermore, G., Schaefers, K., Henry, C., Choi, A., et al. (2025). A phase 3 randomized safety and immunogenicity trial of mRNA-1010 seasonal influenza vaccine in adults. Vaccine 50: 126847. 10.1016/j.vaccine.2025.126847.

10. Moderna, Inc., Moderna announces positive Phase 3 results for seasonal influenza vaccine. Press Release https://feeds.issuerdirect.com/news-release.html?newsid=4899326521164266&symbol=MRNA.

11. Weber, J. S., Carlino, M. S., Khattak, A., Meniawy, T., Ansstas, G., Taylor, M. H., Kim, K. B., McKean, M., Long, G. V., Sullivan, R. J., et al. (2024). Individualised neoantigen therapy mRNA-4157 (V940) plus pembrolizumab versus pembrolizumab monotherapy in resected melanoma (KEYNOTE-942): a randomised, phase 2b study. Lancet 403: 632–644. 10.1016/S0140-6736(23)02268-7.

12. Rojas, L. A., Sethna, Z., Soares, K. C., Olcese, C., Pang, N., Patterson, E., Lihm, J., Ceglia, N., Guasp, P., Chu, A., et al. (2023). Personalized RNA neoantigen vaccines stimulate T cells in pancreatic cancer. Nature 618: 144–150. 10.1038/s41586-023-06063-y.

13. World Health Organization (2022). Evaluation of the quality, safety and efficacy of messenger RNA vaccines for the prevention of infectious diseases: regulatory considerations, Annex 3, TRS No 1039 https://www.who.int/publications/m/item/annex-3-mRNA-vaccines-trs-no-1039.

14. World Health Organization (2005). WHO guidelines on non-clinical evaluation of vaccines, Annex 1, TRS No 927 https://www.who.int/publications/m/item/nonclinical-evaluation-of-vaccines-annex-1-trs-no-927.

15. Li, S. D., and Huang, L. (2008). Pharmacokinetics and biodistribution of nanoparticles. Mol. Pharm. 5: 496–504. 10.1021/mp800049w.

16. European Medicines Agency (2024). Assessment Report. mRESVIA. Moderna. https://www.ema.europa.eu/en/documents/assessment-report/mresvia-epar-public-assessment-report_en.pdf.

17. European Medicines Agency (2021). Assessment Report. COMIRNATY. Pfizer. https://www.ema.europa.eu/en/documents/assessment-report/comirnaty-epar-public-assessment-report_en.pdf.

18. Therapeutic Goods Administration (2021). Nonclinical Summary and Recommendations. SPIKEVAX. Moderna. https://www.tga.gov.au/sites/default/files/2023-07/FOI%204320_0.pdf.

19. Pharmaceuticals and Medical Devices Agency (2021). Report on the Deliberation Results. SPIKEVAX. Moderna. https://www.pmda.go.jp/files/000243267.pdf.

20. Tenforde, M. W., Weber, Z. A., Natarajan, K., Klein, N. P., Kharbanda, A. B., Stenehjem, E., Embi, P. J., Reese, S. E., Naleway, A. L., Grannis, S. J., et al. (2023). Early estimates of bivalent mRNA vaccine effectiveness in preventing COVID-19-associated emergency department or urgent care encounters and hospitalizations among immunocompetent adults - VISION Network, Nine States, September-November 2022. MMWR Morb. Mortal. Wkly Rep. 71: 1637–1646. 10.15585/mmwr.mm7153a1.

21. DeCuir, J., Payne, A. B., Self, W. H., Rowley, E. A. K., Dascomb, K., DeSilva, M. B., Irving, S. A., Grannis, S. J., Ong, T. C., Klein, N. P., et al. (2024). Interim effectiveness of updated 2023-2024 (monovalent XBB.1.5) COVID-19 vaccines against COVID-19-associated emergency department and urgent care encounters and hospitalization among immunocompetent adults aged ≥18 years - VISION and IVY Networks, September 2023-January 2024. MMWR Morb. Mortal. Wkly Rep. 73: 180–188. 10.15585/mmwr.mm7308a5.

22. Link-Gelles, R., Chickery, S., Webber, A., Ong, T. C., Rowley, E. A. K., DeSilva, M. B., Dascomb, K., Irving, S. A., Klein, N. P., Grannis, S. J., et al. (2025). Interim estimates of 2024-2025 COVID-19 vaccine effectiveness among adults aged ≥18 years - VISION and IVY Networks, September 2024-January 2025. MMWR Morb. Mortal. Wkly Rep. 74: 73–82. 10.15585/mmwr.mm7406a1.

23. Surie, D., DeCuir, J., Zhu, Y., Gaglani, M., Ginde, A. A., Douin, D. J., Talbot, H. K., Casey, J. D., Mohr, N. M., Zepeski, A., et al. (2022). Early estimates of bivalent mRNA vaccine effectiveness in preventing COVID-19-associated hospitalization among immunocompetent adults aged ≥65 years - IVY Network, 18 States, September 8-November 30, 2022. MMWR Morb. Mortal. Wkly Rep. 71: 1625–1630. 10.15585/mmwr.mm715152e2.

24. Segal, L., Wouters, S., Morelle, D., Gautier, G., Le Gal, J., Martin, T., Kuper, F., Destexhe, E., Didierlaurent, A. M., and Garçon, N. (2015). Non-clinical safety and biodistribution of AS03-adjuvanted inactivated pandemic influenza vaccines. J. Appl. Toxicol. 35: 1564–1576. 10.1002/jat.3130.

25. Zhang, W., Cui, H., Xu, J., Shi, M., Bian, L., Cui, L., Jiang, C., and Zhang, Y. (2025). Biodistribution and mechanisms of action of MF59 and MF59-like adjuvants. J. Control. Release 378: 573–587. 10.1016/j.jconrel.2024.12.044.

26. Gómez-Mantilla, J. D., Trocóniz, I. F., and Garrido, M. J. (2016). ADME process in vaccines and PK/PD approaches for vaccination optimization. In: Zhou H, TF-P (ed). ADME translational pharmacokinetics/pharmacodynamics of therapeutic proteins: applications in drug discovery and development. John Wiley and Sons, Inc: Hoboken, NJ. pp 1–22. 10.1002/9780470571224.pse558.

27. Marquez-Martinez, S., Salisch, N., Serroyen, J., Zahn, R., and Khan, S. (2024). Peak transgene expression after intramuscular immunization of mice with adenovirus 26-based vector vaccines correlates with transgene-specific adaptive immune responses. PLoS One 19: e0299215. 10.1371/journal.pone.0299215.

28. Reddy, S. T., Rehor, A., Schmoekel, H. G., Hubbell, J. A., and Swartz, M. A. (2006). In vivo targeting of dendritic cells in lymph nodes with poly(propylene sulfide) nanoparticles. J. Control. Release 112: 26–34. 10.1016/j.jconrel.2006.01.006.

29. McCright, J., Naiknavare, R., Yarmovsky, J., and Maisel, K. (2022). Targeting lymphatics for nanoparticle drug delivery. Front. Pharmacol. 13: 887402. 10.3389/fphar.2022.887402.

30. Sebastiani, F., Yanez Arteta, M., Lerche, M., Porcar, L., Lang, C., Bragg, R. A., Elmore, C. S., Krishnamurthy, V. R., Russell, R. A., Darwish, T., et al. (2021). Apolipoprotein E binding drives structural and compositional rearrangement of mRNA-containing lipid nanoparticles. ACS Nano 15: 6709–6722. 10.1021/acsnano.0c10064.

31. Chen, J., Ye, Z., Huang, C., Qiu, M., Song, D., Li, Y., and Xu, Q. (2022). Lipid nanoparticle-mediated lymph node-targeting delivery of mRNA cancer vaccine elicits robust CD8+ T cell response. Proc. Natl. Acad. Sci. U. S. A. 119: e2207841119. 10.1073/pnas.2207841119.

32. Hassett, K. J., Rajlic, I. L., Bahl, K., White, R., Cowens, K., Jacquinet, E., and Burke, K. E. (2024). mRNA vaccine trafficking and resulting protein expression after intramuscular administration. Mol. Ther. Nucleic Acids 35: 102083. 10.1016/j.omtn.2023.102083.

33. Ci, L., Hard, M., Zhang, H., Gandham, S., Hua, S., Wickwire, J., Wehrman, T., Slauter, R., Auerbach, A., Kenney, M., et al. (2023). Biodistribution of Lipid 5, mRNA, and its translated protein following intravenous administration of mRNA-encapsulated lipid nanoparticles in rats. Drug. Metab. Dispos. 51: 813–823. 10.1124/dmd.122.000980.

34. Garneau, N. L., Wilusz, J., and Wilusz, C. J. (2007). The highways and byways of mRNA decay. Nat. Rev. Mol. Cell Biol. 8: 113–126. 10.1038/nrm2104.

35. Wu, D., Chen, Q., Chen, X., Han, F., Chen, Z., and Wang, Y. (2023). The blood–brain barrier: Structure, regulation and drug delivery. Signal Transduct. Target. Ther. 8: 217. 10.1038/s41392-023-01481-w.

36. Shilo, M., Sharon, A., Baranes, K., Motiei, M., Lellouche, J. P., and Popovtzer, R. (2015). The effect of nanoparticle size on the probability to cross the blood-brain barrier: an in-vitro endothelial cell model. J. Nanobiotechnol. 13: 19. 10.1186/s12951-015-0075-7.

37. Betzer, O., Shilo, M., Opochinsky, R., Barnoy, E., Motiei, M., Okun, E., Yadid, G., and Popovtzer, R. (2017). The effect of nanoparticle size on the ability to cross the blood-brain barrier: an in vivo study. Nanomedicine (London) 12: 1533–1546. 10.2217/nnm-2017-0022.

38. Sasson, E., Anzi, S., Bell, B., Yakovian, O., Zorsky, M., Deutsch, U., Engelhardt, B., Sherman, E., Vatine, G., Dzikowski, R., et al. (2021). Nano-scale architecture of blood-brain barrier tight-junctions. eLife 10. 10.7554/eLife.63253.

39. Chander, N., Basha, G., Yan Cheng, M. H., Witzigmann, D., and Cullis, P. R. (2023). Lipid nanoparticle mRNA systems containing high levels of sphingomyelin engender higher protein expression in hepatic and extra-hepatic tissues. Mol. Ther. Methods Clin. Dev. 30: 235–245. 10.1016/j.omtm.2023.06.005.

40. Hou, X., Zaks, T., Langer, R., and Dong, Y. (2021). Lipid nanoparticles for mRNA delivery. Nat. Rev. Mater. 6: 1078–1094. 10.1038/s41578-021-00358-0.

41. Bhattacharjee, B., Lu, P., Monteiro, V. S., Tabachnikova, A., Wang, K., Hooper, W. B., Bastos, V., Greene, K., Sawano, M., Guirgis, C., et al. (2025). Immunological and antigenic signatures associated with chronic illnesses after COVID-19 vaccination. medRxiv: 2025.2002.2018.25322379. 10.1101/2025.02.18.25322379.

42. Patel, A. A., Zhang, Y., Fullerton, J. N., Boelen, L., Rongvaux, A., Maini, A. A., Bigley, V., Flavell, R. A., Gilroy, D. W., Asquith, B., et al. (2017). The fate and lifespan of human monocyte subsets in steady state and systemic inflammation. J. Exp. Med. 214: 1913–1923. 10.1084/jem.20170355.

43. Yang, C., Zhao, H., Espin, E., and Tebbutt, S. J. (2023). Association of SARS-CoV-2 infection and persistence with long COVID. Lancet Respir. Med. 11: 504–506. 10.1016/S2213-2600(23)00142-X.

44. Ogata, A. F., Cheng, C. A., Desjardins, M., Senussi, Y., Sherman, A. C., Powell, M., Novack, L., Von, S., Li, X., Baden, L. R., et al. (2022). Circulating severe acute respiratory syndrome coronavirus 2 (SARS-CoV-2) vaccine antigen detected in the plasma of mRNA-1273 vaccine recipients. Clin. Infect. Dis. 74: 715–718. 10.1093/cid/ciab465.

45. Kent, S. J., Li, S., Amarasena, T. H., Reynaldi, A., Lee, W. S., Leeming, M. G., O’Connor, D. H., Nguyen, J., Kent, H. E., Caruso, F., et al. (2024). Blood distribution of SARS-CoV-2 lipid nanoparticle mRNA vaccine in humans. ACS Nano 18: 27077–27089. 10.1021/acsnano.4c11652.

46. Kowalski, K. G. (1994). An algorithm for estimating the terminal half-life in pharmacokinetic studies. Comput. Methods Programs Biomed. 42: 119–126. 10.1016/0169-2607(94)90048-5.

47. US Food and Drug Administration (2025). Roadmap to reducing animal testing in preclinical safety studies https://www.fda.gov/files/newsroom/published/roadmap_to_reducing_animal_testing_in_preclinical_safety_studies.pdf.

48. US Food and Drug Administration. Inactive ingredient search for approved drug products.

49. International Council for Harmonisation (2023). Nonclinical biodistribution considerations for gene therapy products S12 https://database.ich.org/sites/default/files/ICH_S12_Step4_Guideline_2023_0314.pdf.

50. Boxenbaum, H. (1982). Interspecies scaling, allometry, physiological time, and the ground plan of pharmacokinetics. J. Pharmacokinet. Biopharm. 10: 201–227. 10.1007/BF01062336.

51. Huh, Y., Smith, D. E., and Rose Feng, M. (2011). Interspecies scaling and prediction of human clearance: comparison of small- and macro-molecule drugs. Xenobiotica 41: 972–987. 10.3109/00498254.2011.598582.

52. Huang, Q., and Riviere, J. E. (2014). The application of allometric scaling principles to predict pharmacokinetic parameters across species. Expert Opin. Drug Metab. Toxicol. 10: 1241–1253. 10.1517/17425255.2014.934671.

53. Panther, L., Basnet, S., Fierro, C., Brune, D., Leggett, R., Peterson, J., Pickrell, P., Lin, J., Wu, K., and Lee, H. (2023). 2892. Safety and immunogenicity of mRNA-1647, an mRNA-based cytomegalovirus vaccine in healthy adults: results of a phase 2, randomized, observer-blind, placebo-controlled, dose-finding trial. Open Forum Infect. Dis. 10: ofad500.2475. 10.1093/ofid/ofad500.2475.

54. Moderna Inc. (2022). SPIKEVAX (COVID-19 Vaccine, mRNA) injectable suspension, for intramuscular use. Prescribing information. https://www.fda.gov/media/155675/download.

