## Supplementary material for "Biodistribution of mRNA vaccines in rats: Enrichment in injection site and lymph tissues and rapid clearance without tissue persistence": Goody_Supplement

**Supplemental Material**

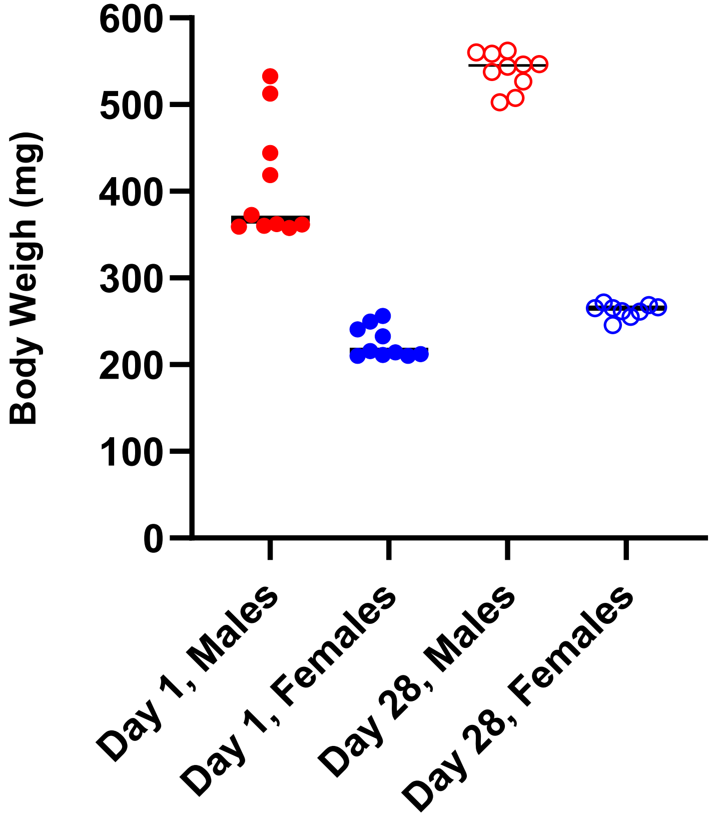


**Supplemental Figure S1. Body weights of male and female rats on Day 1 and Day 28 dosing with mRNA-1273**

Individual body weight values are shown for males (red symbols) and females (blue symbols). n = 8 per group.

### Supplemental Table 1. Summary of mRNA tissue AUC to systemic AUC ratio in male and female Sprague-Dawley rats

| Matrix | | AUC (hr*ng/mL or hr*ng/g) | | | | | | Tissue AUC to systemic AUC ratio (%)[b] | | | | | | |
|---|---|---|---|---|---|---|---|---|---|---|---|---|---|---|
| | mRNA from mRNA-1647[a] | mRNA from NPI-Luc mRNA | | mRNA from mRNA-1273 (original table used hr*pg/mL, converted to hr*ng/mL) | | | | mRNA from mRNA-16 47 | mRNA from NPI-Luc mRNA | | mRNA from mRNA-1273 | | | |
| | Day 1 | Day 1 | | Day 1 | Day 1 | Day 28 | Day 28 | Day 1 | Day 1 | | Day 1 | Day 1 | Day 28 | Day 28 |
| | Male | Male | Female | Male | Female | Male | Female | Male | Male | Female | Male | Female | Male | Female |
| Injection site | 24500 | 26900 | 21000 | 7170 | 51400 | 13200 | 8570 | 101239 | 5434 | 1272 | 2290 | 2050 | 9166 | 556.5 |
| Lymph node (axillary) | ND | NC | 2680 | 5070 | 21300 | 5920 | 23800 | ND | NC | 162 | 1619 | 849 | 4111 | 1545 |
| Lymph node (distal) | 1520 | ND | ND | ND | ND | ND | ND | 6280 | ND | ND | ND | ND | ND | ND |
| Lymph node (inguinal) | ND | NC | NC | ND | ND | ND | ND | ND | NC | NC | ND | ND | ND | ND |
| Lymph node (inguinal/popliteal) | ND | ND | ND | 17100 | 35900 | 12400 | 23600 | ND | ND | ND | 5463 | 1430 | 8611 | 1530 |
| Lymph node (popliteal) | ND | NC | NC | ND | ND | ND | ND | ND | NC | NC | ND | ND | ND | ND |
| Lymph node (proximal) | 4870 | ND | ND | ND | ND | ND | ND | 20123 | ND | ND | ND | ND | ND | ND |
| Spleen | 315 | 26600 | 75400 | 35500 | 107000 | 42900 | 93700 | 1301 | 5373 | 4570 | 11341 | 4262 | 29791 | 6084 |
| **Systemic[b]** | **24.2** | **495** | **1650** | **313** | **2510** | **144** | **1540** | **NA** | **NA** | **NA** | **NA** | **NA** | **NA** | **NA** |
| Bone marrow | 2.70 [c] | 16.4 | 11.7 | ND | ND | ND | ND | 11.15[c] | 3.31 | 0.709 | ND | ND | ND | ND |
| Brain | 0.41[c] | NC | NC | NC | 1.8 | NC | NC | 1.69[c] | NC | NC | NC | 0.0712 | NC | NC |
| Heart | 3.52 [c] | NC | NC | 18.6 | 197 | 12.1 | 276 | 14.5[c] | NC | NC | 5.95 | 7.84 | 8.40 | 17.9 |
| Liver | 11.9 | 90.0 | 508 | 186 | 1630 | 147 | 1050 | 49.2 | 18.2 | 30.8 | 59.4 | 64.9 | 102 | 68.2 |
| Lung | 3.97 | NC[d] | 75.4 | 190 | 387 | 35 | 232 | ND | NC | 4.56 | 60.7 | 15.4 | 24.3 | 15.1 |
| Testes (males) [e] | 4.70 | NC | NA | ND | ND | ND | ND | 19.4 | NC | NC | ND | ND | ND | ND |

Abbreviations: AUC = area under the concentration versus time curve from the start of dose administration to the time after dosing at which the last quantifiable concentration was observed; LNP = lipid nanoparticle; Luc = luciferase; mRNA = messenger RNA; Ratio is expressed in %; Values less than 100% represent tissue exposure less than serum or plasma; NA = not applicable as calculation was not performed because the parameter does not apply to this matrix; NC = not calculable (samples collected but insufficient data points above the limit of quantification for kinetic analysis); ND = not determined as no sample collected per Study Protocol. NPI = nascent peptide imaging.

[a]Average reported across 6 mRNA constructs.

[b]Tissue AUC to plasma/serum AUC ratio (%) for mRNA-1647 study was using plasma where NPI-Luc and mRNA-1273 were using serum.
[c]Only 3 or 4 out of the 6 mRNA constructs were quantifiable for mRNA-1647 Study, results below the limit of quantification were adjusted to a value of zero for the mean calculations.
[d]Concentrations in males were all below LLOQ.
[e]Insufficient ovary tissues from NPI-Luc study to evaluate mRNA.

**Supplemental Table 2. Metabolite profile and identification of Lipid H in rat plasma, urine, and bile following intravenous infusion of Lipid H-containing LNPs[a]**

| Metabolite ID | Retention time (min) | Proposed biotransformation |
|---|---|---|
| M1 | 3.5 | N-Dealkylation- + Ester hydrolysis |
| M2 | 3.9 | Ester hydrolysis (2X) + β-oxidation (2X) |
| M3 | 5.3 | Ester hydrolysis (2X) + β-oxidation (2X) |
| M4 | 6.1 | Ester hydrolysis (2X) |
| M5 | 15.98 | N-Dealkylation- elimination of straight chain ester |
| M6 | 15.94 | Ester hydrolysis (1X) + β-oxidation (1X) |
| M7 | 16.00 | Ester hydrolysis (1X) |
| M8 | 13.92 | Ester hydrolysis (1X) + Aliphatic hydroxylation |
| M9 | 13.75 | Ester hydrolysis (1X) + β-oxidation (1X) + Hydroxylation |
| M10 | 13.83 | Ester hydrolysis (1X) + β-oxidation (1X) + Hydroxylation + Dehydrogenation |
| M11 | 17.85 | Aliphatic hydroxylation + Dehydrogenation |
| M12 | 15.02 | β-oxidation + GSH conjugation |

[a]Lipid H-containing LNPs encapsulated a non-translating mRNA, NT-FIX.

**Supplemental Table 3. Summary of in vitro metabolite profiling of Lipid H[a] in human, rat, and non-human primate hepatocytes**

| Metabolite ID | Retention time | Proposed biotransformation | Human | Rat | NHP |
|---|---|---|---|---|---|
| M1 | 3.2 | N-Dealkylation- + hydrolysis | Y | Y | Y |
| M3 | 6.2 | Ester hydrolysis (2X) + β-oxidation (2X) | Y | N | Y |
| M4 | 6.9 | Ester hydrolysis (2X) | Y | Y | Y |
| M6 | 15.1 | Ester hydrolysis (1X) + β-oxidation (1X) | Y | Y | Y |
| M7 | 16.4 | Ester hydrolysis (1X) | Y | Y | Y |
| Parent Lipid H | 19.2 | NA | Y | Y | Y |

[a]Lipid H-containing LNPs encapsulating a non-translating mRNA, NT-FIX, were used.
Note: Labels for metabolites are consistent with *in vivo* Rat Met ID in Supplemental Table 2 – Y – Detected, N Below Detection limit. M2, M5, M8 to M12 (*in vivo*) were not detected in *in vitro* samples.
